# A new diatom-based multimetric index to assess lake ecological status

**DOI:** 10.1101/2023.03.08.531691

**Authors:** J. Tison-Rosebery, S. Boutry, V. Bertrin, T. Leboucher, S. Morin

## Abstract

Eutrophication impairs lake ecosystems at a global scale. In this context, as benthic microalgae are well-established warnings for a large range of stressors, particularly nutrient enrichment, the Water Framework Directive required the development of diatom-based methods to monitor lake eutrophication.

Here, we present the diatom-based index we developed for French lakes, named IBDL. Data were collected in 93 lakes from 2015 to 2020. A challenge arose from the discontinuous pressure gradient of our dataset, especially the low number of nutrient-impacted lakes. To analyze the data we opted for the so-called “Threshold Indicator Taxa ANalysis” method, which makes it possible to determine a list of “alert taxa”. We obtained a multimetric index based on different pressure gradients (Kjeldahl nitrogen, suspended matter, biological oxygen demand and total phosphorous).

The IBDL proved to be particularly relevant as it has a twofold interest: an excellent relationship with total phosphorus and possible application to any lake metatype. Its complementarity with macrophyte-based indices moreover justifies the use of at least two primary producer components for lake ecological status classification.

## 1 Introduction

Eutrophication is one of the most frequent consequences of human pressure on lake ecosystems at a global scale (Stenger-Kovacs et al., 2007). Primary producers are directly impacted since they are the base of the aquatic food web (Brauer et al., 2012). As the ability of species to compete differs according to nutrient availability, nutrient enrichment results in significant changes in community structure and function (Birk, 2012). For this reason, scientists and policymakers developed indices based on primary producer attributes to monitor eutrophication (Stevenson, 2014). In the early 2000s, the Water Framework Directive (WFD, 2000/06/EC) required all EU member states to implement bioassessment methods based, among other aspects, on the biological quality of “macrophytes and phytobenthos” to assess lake ecological status. This led to the development of numerous methods at the European level.

Poikane et al. (2016) reviewed this panel of methods and observed that countries generally developed separate assessment tools for macrophytes and phytobenthos, and that most of them considered diatoms, which are unicellular microalgae, to be a good proxy for phytobenthos. Diatoms are indeed early and well-established warnings for a large range of stressors, particularly nutrient enrichment (Stevenson, 2014). As a first step, indices originally dedicated to rivers were applied to lakes by the majority of member states (Kelly et al., 2014b), considering that many processes influencing diatom assemblages were comparable between lakeshores and shallow rivers (Cantonati and Lowe, 2014).

In some rare cases, diatom-based indices were developed specifically for lakes, based on species composition and abundance as for rivers (Bennion et al., 2014; Poikane et al., 2016). Diatoms from mud and silts were generally not considered, as they would respond to pore-water chemistry rather than water quality. The recommended sampling substrate varied according to authors, from macrophytes to cobbles or even artificial substrates when no natural substrates are found in all water bodies (King et al., 2006).

To harmonize the different national approaches, a European intercalibration exercise was performed, involving eleven member states (Kelly et al., 2014b). France participated in this exercise with the Biological Diatom Index (BDI, Coste et al., 2009), routinely used to assess river ecological status. Although previous results tended to suggest there was a good correlation between BDI and the environmental pressure gradients, at least in shallow lakes (Cellamare et al., 2012), this intercalibration exercise revealed a poor correlation between BDI values and total phosphorous across France (Kelly et al., 2014b). This was explained by the absence of many lake taxa from the list of key species used to calculate the BDI, resulting in an overall poor relevance of the final status assessment.

The aim of the present study was, therefore, to develop a new diatom-based index for lakes in metropolitan France: the IBDL (*Indice Biologique Diatomées en Lac:* Diatom Biological Index for Lakes). To collect the necessary data, we proposed a method (Morin et al., 2010) consistent with a potential subsequent combination of this index with the existing French macrophyte index IBML (*Indice Biologique Macrophytique en Lac:* Macrophyte Biological Index for Lakes, Boutry et al., 2015). We detail here how diatom data were sampled and analyzed and how we developed the IBDL. Finally, we discuss the relevance of this new index, comparing the results obtained with index scores based on macrophytes, and assessing its ability to reveal environmental gradients.

## 2 Materials and Methods

### 2-1 Data collection

Samples were collected from 93 French lakes (Figure 1) during the summer period, each year from 2015 to 2020, according to Morin et al. (2010). The lakes were classified into three metatypes based on alkalinity, according to the European intercalibration exercise previously performed (Kelly et al., 2014b): low alkalinity (LA, alkalinity ≤ 0.2 meq.1^-1^), medium alkalinity (MA, 0.2 meq.1^-1^ < alkalinity < 1 meq.1^-1^), and high alkalinity (HA, alkalinity ≥ 1 meq.1^-1^). Diatoms were collected from both mineral substrates and lakeshore macrophyte surfaces in observation units (OUs), whose number and location varied according to the lake surface area and the riparian zone types. Such units are defined in the French macrophyte sampling protocol for lakes NF T90-328 (AFNOR, 2022).

**Figure 1:**
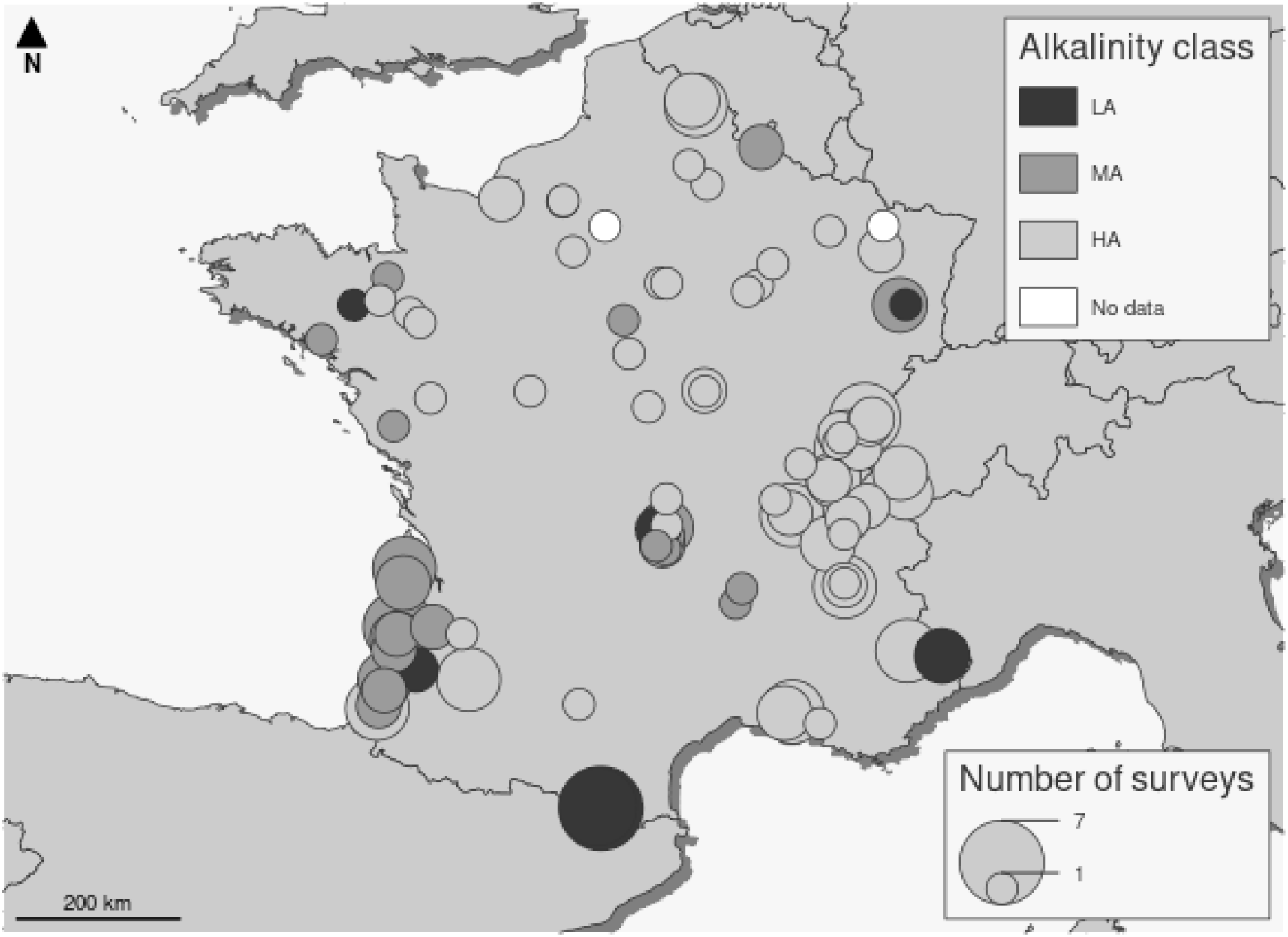
Study sites, number of surveys per site, and lake alkalinity classes (LA: low alkalinity; MA: medium alkalinity; HA: high alkalinity) (Kelly et al., 2014a)

#### 2-1-1 Biological data

Samples from hard mineral substrates were taken from at least five boulders or cobbles selected at random for each OU. The total surface area sampled was equivalent to 100 cm^2^, as defined in the NF T90-354 standard (AFNOR, 2016). Selected substrates had to be submerged within the euphotic zone, at a maximum depth of 0.5 m.

Samples performed on macrophytes were taken from helophytes (mainly *Phragmites australis* (Cav.) Trin. ex Steud.). Green stem segments submerged for at least 4 to 6 weeks were collected from a minimum of 5 macrophytes chosen at random. These stem segments had to be located at a maximum depth of 0.2 m.

Diatoms were sampled from both substrates according to the NF T90-354 protocol, in line with the European standards (EN 13946, European Commission). Cells were identified at 100x magnification by examining permanent slides of cleaned diatom frustules (400 valves per slide) using, among others, Krammer and Lange-Bertalot (1986-1991) and Lange-Bertalot (1995–2015, 2000–2013). Taxonomic homogenization was performed with Omnidia 6 software (Lecointe et al., 1993).

All OUs from a single lake were sampled within a maximum of 21 days. Diatom counts had to include at least 350 cells per slide, with more than 50% of the diatom cells determined at the species level, to comply with the NF T90-354 requirements.

#### 2-1-2 Physico-chemical data

Parameter values were determined in summer at the deepest point of each lake, according to European standards. Data were obtained from national surveillance monitoring programs. Water quality analysis was not systematically performed each year: in a few cases, the most recent physicochemical data available were collected three years before the diatom samples. The following parameters were recorded: biological oxygen demand (BOD5, mg.1^-1^), oxygen (O_2_, mg.1^-1^), oxygen saturation (% O_2_), conductivity (Cond, μs.cm^2^), Kjeldahl nitrogen (NKJ, mg.1^-1^), ammonium (NH_4_, mg.1^-1^), nitrates (NO_3_, mg.1^-1^), nitrites (NO_2_, mg.1^-1^), orthophosphates (PO_4_, mg.1^-1^), total phosphorous (Pt, mg.1^-1^) and suspended particles (SP, mg.1^-1^).

### 2-2 Data analysis and index settlement

All analyses were performed with R version 4.1.2 (2021-11-01) (R Core Team, 2021) (Platform: x86_64-pc-linux-gnu (64-bit), Running under: Ubuntu 22.04.1 LTS).

Considering that the final dataset revealed a discontinuous trophic gradient, we opted for the so-called “Threshold Indicator Taxa ANalysis” method (TITAN2 package, Baker et al., 2020), which, based on bootstrapping and permutations, makes it possible to determine a list of “alert taxa”. The presence and/or increasing abundance of alert taxa reveal the existence of anthropogenic pressures. TITAN replaces the community□ level response along a composite gradient with taxon □ specific responses towards single environmental variables (Dufrêne and Legendre 1997). Negative and positive responses are distinguished, and cumulative decreasing or increasing responses in the community are tracked. This method is particularly suitable for setting up multimetric indices.

A three-step procedure was necessary to build our Biological Diatom Index for Lakes (IBDL): identification of alert taxa, choice of relevant metrics, and aggregation of these metrics to obtain the final index score.

#### 2-2-1 Identification of alert taxa

For the next part of the analysis, we set an occurrence threshold ≥3 for taxa to be included in the index calculation (the so-called “index taxa”).

TITAN combines change-point analysis (nCPA; King and Richardson 2003) and indicator species analysis (IndVal, Dufrêne and Legendre 1997). Basically, the change-point analysis compares within-group vs. between-group dissimilarity to detect shifts in community structure along the environmental variable considered (for further details concerning this method see Baker and King, 2010). Indicator species analysis then identifies the strength of association between any particular taxon and this sample grouping. At the end of the process, two IndVal scores are calculated for a single taxon in a two-group classification. The algorithm finally classifies taxa into three different categories: Z^+^ taxa, showing a significant increase in abundance along the increasing environmental gradient; Z^-^ taxa, showing a significant decrease along this gradient; and indifferent taxa, with no significant trend.

Alert taxa were defined as Z^+^ or Z^-^ taxa whose shift thresholds were greater or lesser than the community shift threshold.

#### 2-2-2 Building metrics and selecting the relevant ones

For each environmental variable, a metric was calculated at the OU scale according to (1):

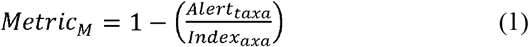

Where:

Alerttaxa is the number of alert taxa and Indextaxa is the number of index taxa in the sample.

The metric value is bounded between 0 and 1. The lowest value (0) corresponds to a species list entirely composed of alert taxa (determined for the environmental variable considered).

To build our index, we then selected the most relevant metrics, i.e., those with the best relationship with the environmental parameter considered. We used Pearson’s correlation coefficients to measure this statistical association and only kept metrics showing a Pearson’s coefficient over 0.6. Metrics should significantly increase with impairment, significantly decrease with impairment, or show no particular pattern. We obtained the response patterns of the different metrics by transforming raw values into normalized deviations (Standardized Effect Size: SES, Gotelli and McCabe, 2002; Mondy et al., 2012) (2). SES values made it possible to obtain a single response pattern for a metric whatever the lake metatype and substrate type considered.

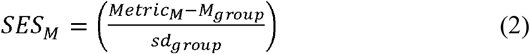

Where:

Metric_M_ is the observed value of the metric, M_group_ and sd_group_ are the mean and standard deviation, respectively, of the metric value for a given group of samples (i.e., substrate type x lake alkalinity metatype) (values of M_group_ and sd_group_ are given in Table 1 S1)

The next step consisted of the normalization of SES values (SESnor_M_) to make comparable metric variation ranges (3):

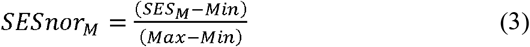

Where:

SES_M_ is the observed value of SES for a given metric, Min its minimum value and Max its maximum value in the whole dataset (values of Min and Max are given in Table 2 S1).

We further transformed metric values from normalized SES into the Ecological Quality Ratio (EQR) (4), i.e. the ratio between the observed value of a metric (SESnor_M_) and its expected value under reference conditions (Kelly et al., 2014a), for any lake metatype and any substrate (SESnor_Mref_, values given in Table 3 S1).

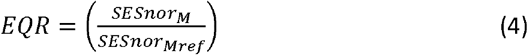

Finally, for each metric, we performed a Wilcoxon test to detect the potential influence of substrate type on the EQR values obtained at the OU scale.

#### 2-2-4 Aggregating metric values to obtain the final IBDL score

The final index score was obtained at the OU scale by averaging the selected metric values, expressed in EQR.

For a score calculated for both mineral and macrophyte substrates, the lowest value was considered the final score.

Each OU belongs to one of the four riparian zone types, as required in the NF T90-328 standard (AFNOR, 2022). These types were defined from the vegetation composition and/or anthropogenic alterations of the lakeshore. The percentage of each riparian zone type was estimated *in situ*, on the whole lake perimeter, during the sampling surveys. The final index score for the whole lake was derived from a weighted average of the ScoreOU (5), taking into account the percentage of the lake perimeter each OU represented in terms of riparian zone type (Pctype).

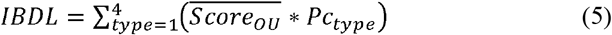

Finally, the resulting IBDL scores varied between 0 (worst water quality) and 1. Relationships between IBDL scores and the different environmental variables considered were tested *a posteriori* with simple linear regressions (R “mass” package, Venables & Ripley, 2002).

### 2-3 Comparing IBDL and IBML scores

We compared IBDL and IBML scores, based respectively on diatom and macrophyte communities, to evaluate their complementarity or redundancy. IBML scores were computed with the online application https://seee.eaufrance.fr/api/indicateurs/IBML/1.0.1 and the “httr” package (Wickham, 2022).

We built a multiple linear regression model (“mass” package) to test which index correlated best with Pt values: IBML, IBDL or a combination of both (mean value).

### 2-4 Preparing intercalibration

Considering a future intercalibration exercise, we analyzed the relationships between IBDL scores and Pt for each lake metatype. A good correlation of the candidate metric with Pt constitutes a key criterion for considering the index ready for integration into the intercalibration process (Kelly et al., 2014b).

We also plotted IBDL against CM scores (intercalibration Common Metric, i.e., the Trophic Index developed by Rott et al., 1998), to check their compliance. The CM was calculated with Omnidia 6 software.

## 3- Results

Our data revealed discontinuous pressure gradients (Table 1), with a clear lack of impacted conditions and an over-representation of lakes characterized by low eutrophication levels.

**Table 1.** Physico-chemical data available for analysis. sd: standard deviation, p25: 25^th^ percentile; p75: 75^th^ percentile.

| Variable | % of missing values | mean | sd | median | p25 | p75 | maximum |
| --- | --- | --- | --- | --- | --- | --- | --- |
| ammonium (NH<sub>4</sub>, mg.l-1) | 0.292 | 0.090 | 0.350 | 0.015 | 0.010 | 0.060 | 3.30 |
| biological oxygen demand (DBO<sub>5</sub>, mg.l-1) | 0.584 | 2.157 | 2.615 | 1.300 | 0.900 | 1.800 | 12.00 |
| conductivity | 0.309 | 230.108 | 124.368 | 243.500 | 158.000 | 297.000 | 815.00 |
| Kjeldahl nitrogen (NKJ, mg.l-1) | 0.292 | 0.661 | 0.959 | 0.250 | 0.250 | 0.700 | 6.90 |
| nitrates (NO<sub>3</sub>, mg.l-1) | 0.292 | 1.113 | 1.222 | 0.600 | 0.250 | 1.400 | 6.07 |
| nitrites (NO<sub>2</sub>, mg.l-1) | 0.292 | 0.011 | 0.025 | 0.005 | 0.005 | 0.010 | 0.30 |
| orthophosphates (PO<sub>4</sub>, mg.l-1) | 0.292 | 0.015 | 0.026 | 0.005 | 0.005 | 0.010 | 0.22 |
| oxygen (O<sub>2</sub>, mg.l-1) | 0.333 | 9.143 | 3.943 | 8.700 | 8.100 | 9.685 | 73.00 |
| oxygen saturation (% O<sub>2</sub>) | 0.333 | 110.203 | 20.910 | 108.000 | 101.000 | 117.650 | 187.00 |
| suspended particles (SP, mg.l-1) | 0.292 | 7.979 | 18.145 | 2.800 | 1.600 | 5.000 | 153.00 |
| total phosphorous (Pt, mg.l-1) | 0.292 | 0.027 | 0.067 | 0.005 | 0.005 | 0.015 | 0.51 |

Biotic and abiotic data were obtained for 958 samples. Considering the data validation criteria, 99% of the samples were included in the analysis. Data from both substrate types were available for 552 OUs. Seven hundred eighty taxa were recorded, 8% of which were identified to the genus level. One hundred and twenty-one alert taxa were determined out of 590 index taxa (S2).

We obtained the following Pearson test values for the different metrics at the OU scale: R = −0.715 for the metric based on the parameter NKJ, R = −0.754 for BOD_5_, R = −0.688 for Pt, R = −0.666 for SP, R = −0.553 for PO_4_, R = −0.329 for Conductivity, R = −0.174 for O_2_, R = −0.265 for NO_2_, and R = −0.204 for %O_2_. Considering the selection rule proposed (|R|>0.6), only the metrics based on NKJ, BOD5, Pt and SP were considered to build the IBDL.

Metric values (in EQR) calculated from the lists of taxa sampled on mineral substrates and macrophytes for a single OU did not differ significantly (p-value =0.65).

IBDL scores at the lake level were calculated from the selected metrics following the aggregation rules proposed. The scores obtained were distributed as given in Figure 2. IBDL could not be calculated for 20% of the samples due to incomplete floristic data.

**Figure 2:**
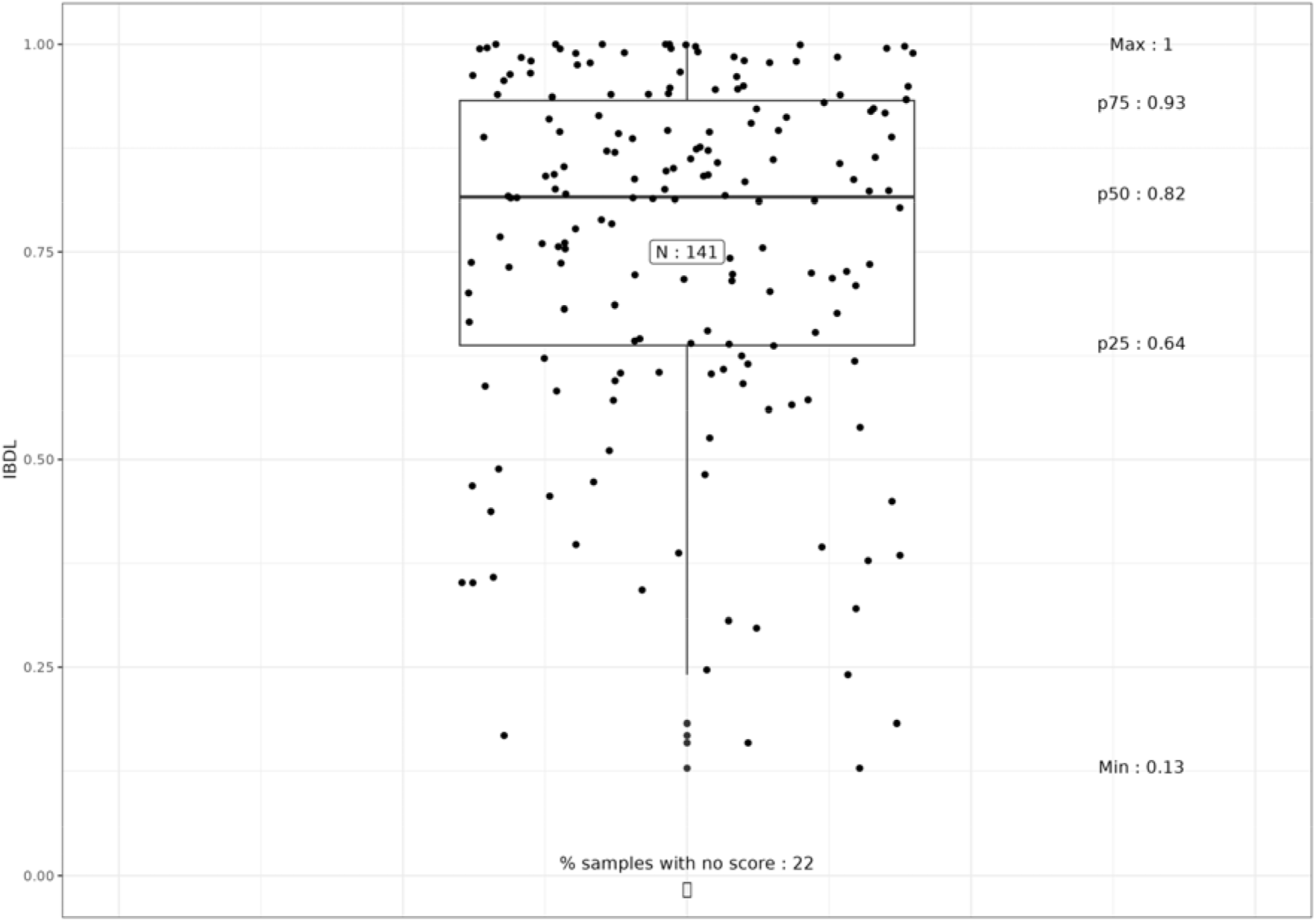
Distribution of the IBDL scores obtained (p25: 25^th^ percentile; p50: median value; p75: 75^th^ percentile).

The relationships between IBDL scores and the different environmental variables considered were very good (Figure 3) in both high-alkalinity and medium-alkalinity lakes. IBDL scores showed high correlations with these variables, particularly Pt, in both high alkalinity (R^2^ = 0.63, p = 1.8e^-15^) and medium alkalinity lakes (R2 = 0.83, p = 8.3e^-11^). Note that data from low alkalinity lakes were too scarce to perform such correlations.

**Figure 3.**
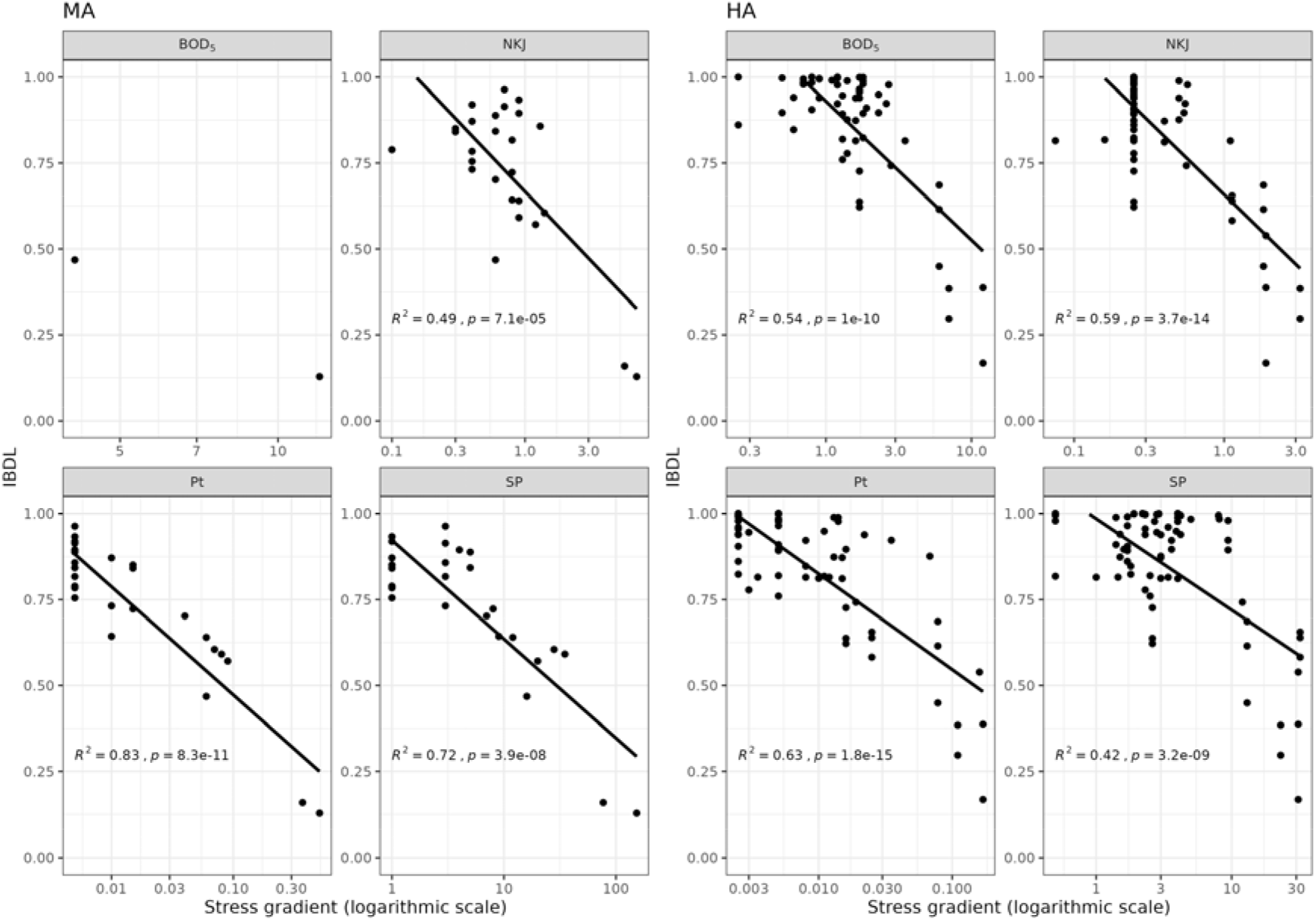
Relationships between IBDL and the environmental variables considered (MA: medium-alkalinity lakes; HA: high-alkalinity lakes; BOD_5_: biological oxygen demand; NKJ: Kjeldahl nitrogen; Pt: total phosphorous; SP: suspended particles.

IBDL scores were also strongly associated with CM scores (R2 = 0.52 and p = 2.2e^-16^ for high-alkalinity lakes; R2 = 0.87 and p = 1.8 e^-7^ for medium-alkalinity lakes) (Figure 4).

**Figure 4:**
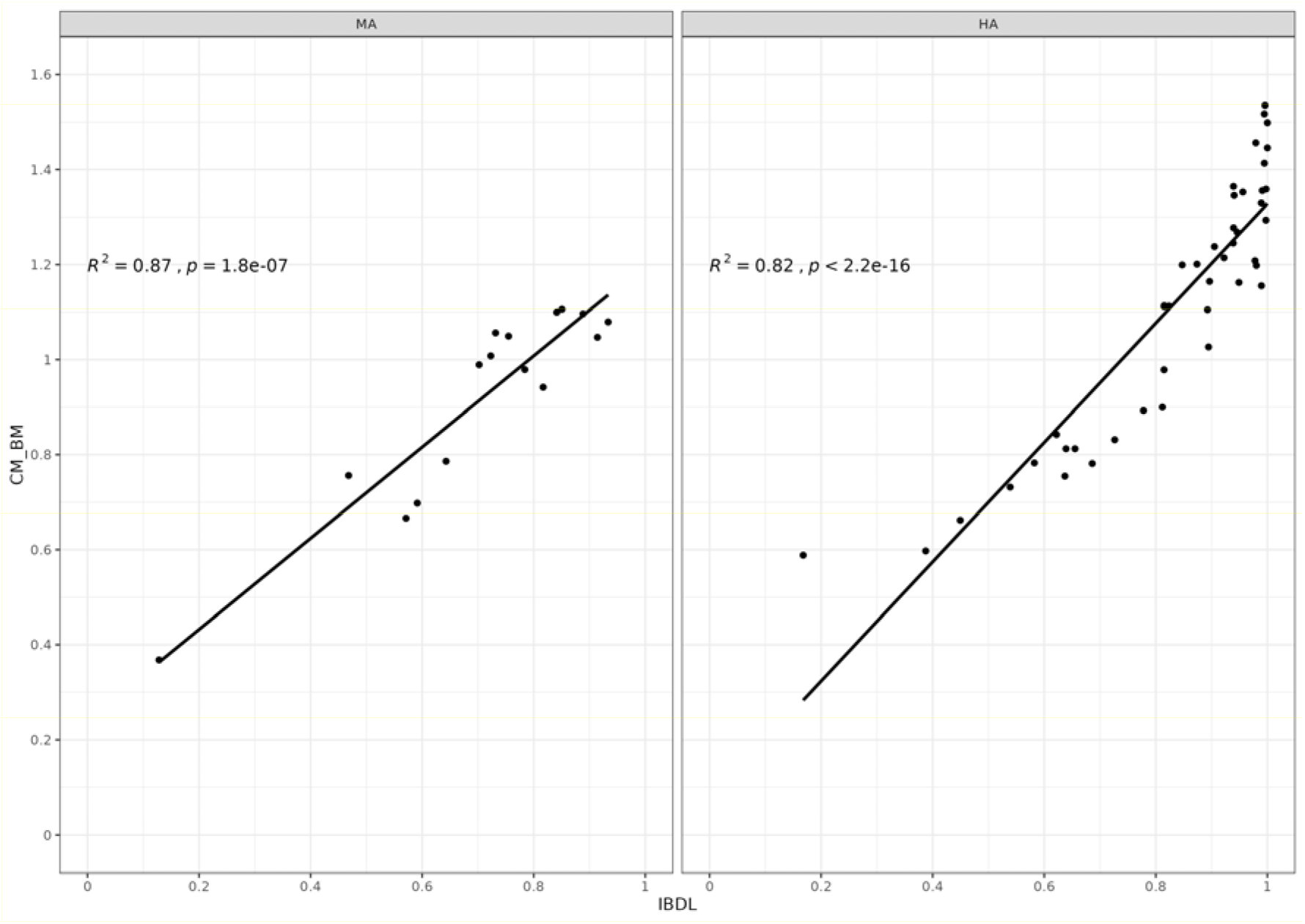
Relationships between IBDL and the common metric (CM) in medium alkalinity (MA) and high alkalinity (HA) lakes.

IBDL scores showed a better correlation with Pt (AIC = −171.44) than did IBML (AIC = −129.25) or a combination of both indices (AIC = −169.44). Nevertheless, IBDL tended to be generally less stringent than IBML (in 18 out of 22 samples), especially for scores higher than 0.8 (clearly dominant here). Figure 5 presents the difference between IBDL and IBML scores according to IBDL scores.

**Figure 5:**
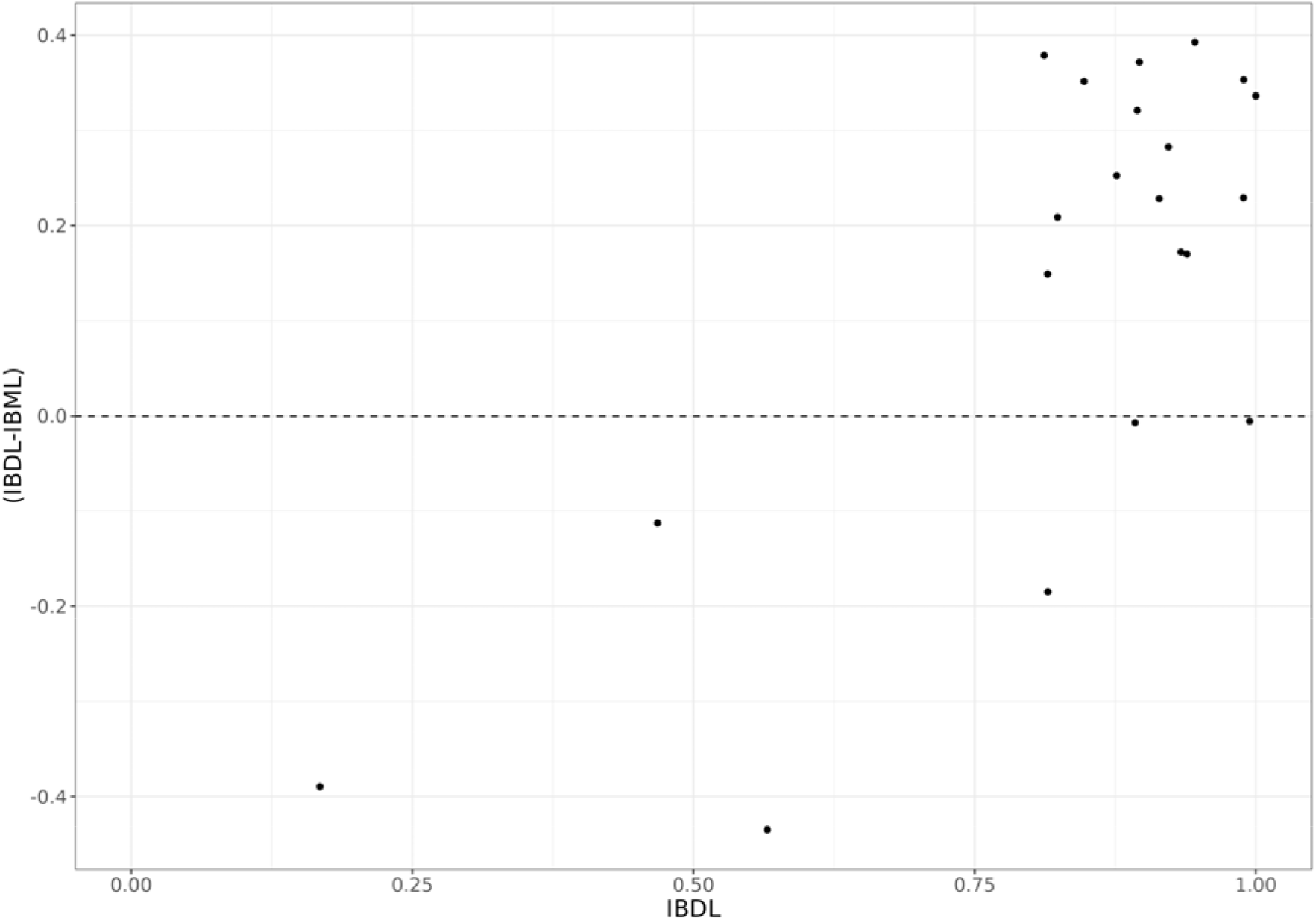
Difference between IBDL and IBML scores according to IBDL scores

## 4 Discussion

As required by the WFD, we developed a diatom index for the assessment of the ecological status of French lakes. We obtained very good correlations between IBDL and key environmental variables. One major challenge arose from the discontinuous pressure gradient of our dataset, especially the low available number of nutrient-impacted lakes.

The scarcity of impacted lakes in the datasets used to build diatom indices is not rare and has already been pointed out by some authors (Bennion et al., 2014). This lack makes it impossible to capture the entire trophic gradient or to build reliable species’ ecological profiles. However, the majority of existing indices are calculated as an abundance-weighted average of the ecological profiles of every taxon from a sample, according to the Zelinka & Marvan formula (Zelinka and Marvan, 1961). This method is far from optimal for datasets showing discontinuous or very specific environmental conditions (Carayon et al., 2020). In such cases, the identification of alert taxa seems more appropriate than considering diatom communities as a whole. This has made the TITAN algorithm increasingly popular for detecting specific taxa providing reliable signals of a specific stress (Khamis et al., 2014; Costas et al., 2018; Gieswein et al., 2019; Carayon et al., 2020; Gonzalez-Paz et al., 2020).

Using this method, we built a multimetric index based on different pressure gradients (NKJ, SP, BOD5 and Pt). Although the strong influence of nutrients and organic matter on diatom community composition is well established (Jüttner, 2010; Stevenson et al., 2013), diatombased metrics rarely take into account suspended particles for water quality assessment (but see Larras et al., 2017). Diatoms are indeed directly impaired by turbidity, reducing light availability for photosynthesis. Multimetric indices thus offer simple tools to summarize the effect of multi-pressure gradients on communities (Riato et al., 2018), and can be considered more effective for assessing biological conditions than a single metric (Stevenson et al., 2013). However, despite their increasing use, multimetric indices suffer from the subjectivity that can arise from metric selection (Reavie et al., 2008). Here, we attempted to avoid this pitfall by proposing a method of selecting metrics based on the robustness of their response to environmental gradients.

IBDL appears less stringent than IBML when assessing lakes’ ecological status. Literature comparing results from different indices in lakes, though scarce, tends to agree with this overestimation of water quality by diatom-based methods (Kolada et al., 2016). Phytobenthos has long been paid less attention than macrophytes for the assessment of lake ecological status. It is true that recent diatom-based metrics barely detected newly impacted lakes that would not have been detected by macrophyte metrics. Bennion et al. (2014) showed, for example, that their index (LTDI) performed well for lakes with good ecological status, but diatoms and other methods agreed less for lakes of lower status. This was particularly the case in the presence of morphological alterations, for which diatoms are poor indicators. A possible general explanation for the lower stringency of diatom-based indices in lakes is the high abundance of species complexes like *Achnanthidium minutissimum* or *Gomphonema parvulum*. Such complexes merge taxa that are morphologically close but with different ecological preferences. Due to the existence of different taxa within the *A. minutissimum* complex, many authors consider it an indicator of good water quality (Almeida et al., 2014), whereas others consider it as tolerant towards toxic contaminants (micropollutants) and hydrologic disturbances (Cantonati et al., 2014; Lainé et al., 2014). Considering the generally high abundance of *A. minutissimum* in samples, this tends to blur the overall pressure-response relationship between index scores and environmental variables (Potapova and Hamilton, 2007). TITAN provides a means to avoid this pitfall, as such complexes are not selected as alert taxa, given that their abundance dynamics do not show clear response patterns to environmental gradients. Indeed, *A. minutissimum*, although highly abundant in our dataset (22% of total species abundances), was not considered an alert taxon. The fact remains that IBDL tends to be less stringent than IBML, despite better relationships with Pt. In consequence, we have to explain why we think that the use of diatom-based indices to assess lake ecological status is justified.

First, the discrepancy between macrophyte and diatom responses relies mainly on the differences between their integration periods, given that indices provide information on ecological conditions over the time an assemblage develops. Lavoie (2009) showed the integration period of diatom-based indices to be about 2–5 weeks for nutrients, whereas macrophytes react on yearly time scales (Kelly et al., 2016). As diatoms catch nutrients directly from the water column (Wetzel, 2001), they also may be more directly sensitive to rapid changes in trophic status than macrophytes (Vermaat et al., 2022). The rapid response of phytobenthos should justify its routine use (Schneider et al., 2019), in particular for lakes in non-equilibrium states (Kelly et al., 2016).

Second, diatom-based indices are essential where hydrologic pressures in littoral areas prevent the development of macrophytes, and in lake typologies where macrophyte communities are naturally species-poor or even absent (Schneider et al., 2019). Thus, while macrophyte-based indices cannot be calculated in all lakes, this is not true for diatom-based indices. Moreover, our results show that, with IBDL, water quality managers can directly compare ecological status assessments from different lakes even if the substrate sampled is different. Many studies highlighted that allelopathic relationships between macrophytes and epiphytic diatoms may be responsible for specific associations between macrophytes and diatom species and, thus, may contribute to the organization of particular assembly patterns (Hinojosa-Garro et al., 2010). In any case, in terms of ecological preferences, and consequently in terms of IBDL scores, our results did not show any significant differences between communities sampled on mineral substrates or macrophytes at the OU level, corroborating previous results obtained by Kitner and Poulíčková (2003) and Bennion et al. (2014). Other studies even support the use of epiphytic diatoms as biological indicators for lakes irrespective of the dominant macrophyte species sampled (Cejudo-Figueiras et al., 2010). The key point is to avoid senescent material or recently grown shoots that would potentially induce a colonization stage effect (King et al., 2006).

The next challenge was to check the consistency of the resulting classification of lakes based on IBDL to the harmonized definition of good ecological status established in the completed intercalibration exercise (Kelly et al., 2014b). The first step consisted in testing the correlation between IBDL scores and total phosphorus in our dataset. Only HA and MA typologies were considered here but, in any case, the last intercalibration exercise could not be performed for LA lakes. We obtained very good correlations that are clearly an improvement compared to the non-significant relationship previously obtained between BDI (diatom index used for the assessment of rivers) and Pt, and even better than the pressure-impact relationships observed at a pan-European scale (R^2^ between national methods and Pt ranged from 0.32 to 0.66 max., Kelly et al., 2014b). The second step consisted in testing the correlation between IBDL scores and the intercalibration common metric (CM) scores, in EQR. Here, the correlations demonstrated a very good agreement between IBDL and CM scores in both medium (R2 = 0.87) and high alkalinity (R2 = 0.82) lakes. We are, therefore, confident in our ability to match IBDL ecological status thresholds with those validated at the European level.

## Conclusion

The new diatom index proposed here meets the requirements of the WFD and makes it possible to assess lakes’ ecological status in metropolitan France. The IBDL has proved to be particularly relevant as it has a twofold interest: an excellent relationship with total phosphorus and an application in any lake metatype. Its complementarity with IBML justifies the use of at least two primary producer components for ecological status classification (Kelly et al., 2016).

## Acknowledgments

We thank all Water Agencies for data sharing, all Regional Departments for Environment for data collection, and the French Biodiversity Agency (OFB, pôle ECLA) for its financial support. We also thank the two reviewers for their helpful comments on this work.

## Author’s contributions

All authors participated in designing the study and developing aims and research questions. S.B. designed methodology, extracted data and made the analyses, supported by T.L. concerning pretreatments before intercalibration. J.T.R. led the writing of the manuscript supported by S.B., S.M. and V.B. All authors contributed critically to the drafts, contributed to the final version of the manuscript, and gave final approval for publication.

