## Supplementary material 1 Table 3 for "A new diatom-based multimetric index to assess lake ecological status"

| variable | SESnorRef |
|---|---|
| BOD5 | 0.9358017 |
| MES | 0.9081657 |
| NKJ | 0.9138570 |
| NO2 | 0.8865007 |
| NO3 | 0.7155957 |
| PO4 | 0.9088790 |
| Pt | 0.8880905 |
| cond__invivo | 0.7861347 |
| o2_dissous__invivo | 0.9069739 |
| sat_o2__invivo | 0.9519387 |
