## Supplementary material 2 for "A new diatom-based multimetric index to assess lake ecological status"

| code | name | author | Alert taxa (1: yes ; O: no) BOD5 | SP | NKJ | Pt |
|---|---|---|---|---|---|---|
| AAMB | *Aulacoseira ambigua* | (Grunow) Simonsen | 1 | 0 | 1 | 1 |
| ABRT | *Achnanthidium bioretii* | (Germain) Edlund | 0 | 0 | 0 | 0 |
| ABRT | *Achnanthidium bioretii* | (Germain) Monnier Lange-Bertalot & Ector | 0 | 0 | 0 | 0 |
| ABRY | *Adlafia bryophila* | (Petersen) Lange-Bertalot in Moser & al. | 0 | 0 | 0 | 0 |
| ACAF | *Achnanthidium affine* | (Grun) Czarnecki | 0 | 0 | 0 | 0 |
| ACLI | *Achnanthidium lineare* | W. Smith | 0 | 0 | 0 | 0 |
| ACOP | *Amphora copulata* | (Kützing) Schoeman & Archibald | 1 | 1 | 1 | 0 |
| ADAM | *Achnanthidium atomoides* | Monnier, Lange-Bertalot & Ector | 0 | 0 | 0 | 0 |
| ADAS | *Achnanthidium anastasiae* | (Kaczmarska) Chudaev et Gololobova | 0 | 0 | 0 | 0 |
| ADCA | *Achnanthidium caledonicum* | Lange-Bertalot )Lange-Bertalot | 0 | 0 | 0 | 0 |
| ADCT | *Achnanthidium catenatum* | (Bily & Marvan) Lange-Bertalot | 0 | 0 | 0 | 1 |
| ADDA | *Achnanthidium daonense* | (Lange-Bertalot) Lange-Bertalot Monnier & Ector | 0 | 0 | 0 | 0 |
| ADEG | *Achnanthidium exiguum* | (Grunow) Czarnecki | 1 | 0 | 0 | 1 |
| ADEU | *Achnanthidium eutrophilum* | (Lange-Bertalot)Lange-Bertalot | 1 | 1 | 0 | 0 |
| ADEX | *Achnanthidium exile* | (Kützing) Heiberg | 0 | 0 | 0 | 0 |
| ADEX | *Achnanthidium exile* | (Kützing) Bukhtiyarova | 0 | 0 | 0 | 0 |
| ADEX | *Achnanthidium exile* | (Kützing) Round & Bukhtiyarova | 0 | 0 | 0 | 0 |
| ADGL | *Achnanthidium gracillimum* | (Meister)Lange-Bertalot | 0 | 0 | 0 | 0 |
| ADHE | *Achnanthidium helveticum* | (Hustedt) Monnier Lange-Bertalot | 0 | 0 | 0 | 0 |

| | | | Alert taxa (1: yes ; 0: no) | | | |
|---|---|---|---|---|---|---|
| code | name | author | BOD5 | SP | NKJ | Pt |
| | | & Ector | | | | |
| ADJK | *Achnanthidium jackii* | Rabenhorst | 0 | 0 | 0 | 0 |
| ADKR | *Achnanthidium kranzii* | (Lange-Bertalot) Round & Bukhtiyarova | 0 | 0 | 0 | 0 |
| ADLA | *Achnanthidium latecephalum* | Kobayasi | 0 | 0 | 0 | 0 |
| ADMC | *Achnanthidium microcephalum* | Kützing sensu W. Smith | 0 | 0 | 0 | 0 |
| ADMI | *Achnanthidium minutissimum* | (Kützing) Czarnecki group 1 | 0 | 0 | 0 | 0 |
| ADMI | *Achnanthidium minutissimum* | (Kützing) Czarnecki group 2 | 0 | 0 | 0 | 0 |
| ADMI | *Achnanthidium minutissimum* | (Kützing) Czarnecki group 3 | 0 | 0 | 0 | 0 |
| ADMI | *Achnanthidium minutissimum* | (Kützing) Czarnecki | 0 | 0 | 0 | 0 |
| ADMO | *Achnanthidium delmontii* | PeRes. Le Cohu et Barthes | 0 | 0 | 0 | 0 |
| ADMS | *Adlafia minuscula* | (Grunow) Lange-Bertalot | 0 | 0 | 0 | 0 |
| ADMU | *Adlafia muralis* | (Grunow in Van Heurck 1880) Li et Qi | 0 | 0 | 0 | 0 |
| ADMU | *Adlafia muralis* | (Grunow) Monnier & Ector | 0 | 0 | 0 | 0 |
| ADNM | *Achnanthidium neomicrocephalum* | Lange-Bertalot & Staab | 0 | 0 | 0 | 0 |
| ADPL | *Achnanthidium pseudolineare* | Van de Vijver. Novais et Ector | 0 | 0 | 0 | 0 |
| ADPS | *Achnanthidium petersenii* | (Hustedt) C.E. Wetzel,Ector, D.M. Williams & Jüttner | 0 | 0 | 0 | 0 |
| ADPY | *Achnanthidium pyrenaicum* | (Hustedt) Kobayasi | 0 | 0 | 0 | 0 |
| ADRI | *Achnanthidium rivulare* | Potapova & Ponader | 0 | 0 | 0 | 1 |
| ADRK | *Achnanthidium rosenstockii* | (Lange-Bertalot) Lange-Bertalot in Krammer & Lange-Bertalot | 0 | 0 | 0 | 0 |
| ADRU | *Achnanthidium druartii* | Rimet & Couté in Rimet & al. | 0 | 1 | 0 | 0 |

| code | name | author | Alert taxa (1: yes ; O: no) | | | |
|---|---|---|---|---|---|---|
| | | | BOD5 | SP | NKJ | Pt |
| ADSA | *Achnanthidium saprophilum* | (Kobayasi et Mayama) Round & Bukhtiyarova | 0 | 0 | 0 | 0 |
| ADSB | *Achnanthidium straubianum* | (Lange-Bertalot)Lange-Bertalot | 0 | 0 | 0 | 0 |
| ADSH | *Achnanthidium subhudsonis* | (Hustedt) H. Kobayasi | 0 | 0 | 0 | 0 |
| ADSO | *Achnanthidium subatomoides* | (Hustedt) Monnier, Lange-Bertalot et Ector | 0 | 0 | 0 | 0 |
| ADSU | *Achnanthidium subatomus* | (Hustedt) Lange-Bertalot | 0 | 0 | 0 | 0 |
| ADTC | *Achnanthidium tropicocatenatum* | Marquardt, C.E.Wetzel & Ector | 0 | 0 | 0 | 0 |
| ADTR | *Achnanthidium trinode* | Ralfs in Pritchard | 0 | 0 | 0 | 0 |
| AFOR | *Asterionella formosa* | Hassall | 0 | 0 | 0 | 0 |
| AFOR | *Asterionella formosa* | Hassall | 0 | 0 | 0 | 0 |
| AGRU | *Achnanthes grubei* | Simonsen | 0 | 0 | 0 | 0 |
| AGSL | *Aulacoseira granulata* | (Ehrenberg) Simonsen | 1 | 0 | 1 | 1 |
| AGSL | *Aulacoseira granulata* | (Ehrenberg) Simonsen | 1 | 0 | 1 | 1 |
| AHOF | *Achnanthidium hoffmannii* | Van de Vijver. Ector, Mertens & Jarlman | 0 | 0 | 0 | 0 |
| AINA | *Amphora inariensis* | Krammer | 0 | 0 | 0 | 0 |
| ALBL | *Adlafia langebertalotii* | Monnier et Ector | 0 | 0 | 0 | 0 |
| AMCD | *Amphora macedoniensis* | Nagumo | 0 | 0 | 0 | 0 |
| AMDN | *Amphora meridionalis* | Levkov | 0 | 0 | 0 | 0 |
| AMID | *Amphora indistincta* | Levkov | 0 | 0 | 0 | 0 |
| AMLB | *Amphora lange-bertalotii* | Levkov, & Metzeltin | 1 | 1 | 1 | 1 |
| AMUZ | *Aulacoseira muzzanensis* | (Meister) Krammer | 1 | 1 | 1 | 1 |
| ANMN* | *Actinocyclus* | (Gregory ex | 0 | 0 | 0 | 0 |

| code | name | author | Alert taxa (1: yes ; O: no) | | | |
|---|---|---|---|---|---|---|
| | | | BOD5 | SP | NKJ | Pt |
| | *normanii morphotype normanii* | Greville) Hustedt | | | | |
| ANRS | *Aneumastus rosettae* | Lange-Bertalot & Miho | 0 | 0 | 0 | 0 |
| ANSS | *Aneumastus stroesei* | (Østrup) Mann & Stickle in Round Crawford & Mann | 0 | 0 | 0 | 0 |
| AOVA* | *Amphora ovalis var. ovalis* | (Kützing) Kützing | 0 | 0 | 0 | 0 |
| APED | *Amphora pediculus* | (Kützing) Grunow | 0 | 1 | 0 | 1 |
| APEL | *Amphipleura pellucida* | Kützing | 0 | 0 | 0 | 0 |
| APFI | *Achnanthidium pfisteri* | Lange-Bertalot | 0 | 0 | 0 | 0 |
| AUAL | *Aulacoseira alpigena* | Grunow) Krammer | 0 | 0 | 0 | 0 |
| AUGR | *Aulacoseira granulata* | (Ehrenberg) Simonsen | 1 | 1 | 1 | 1 |
| AUGR | *Aulacoseira granulata* | (Ehrenberg) Simonsen | 1 | 1 | 1 | 1 |
| AUPU | *Aulacoseira pusilla* | (Meister) Tuji et Houki | 1 | 1 | 1 | 1 |
| AUSL | *Aulacoseira scalaris* | (Grunow) Houk, Klee & Passauer | 0 | 0 | 0 | 0 |
| AUSU | *Aulacoseira subarctica* | (O. Müller) Haworth | 0 | 0 | 0 | 0 |
| AUTL | *Aulacoseira tenella* | (Nygaard) Simonsen | 0 | 0 | 0 | 0 |
| AUVA | *Aulacoseira valida* | Grunow)Krammer | 0 | 1 | 1 | 1 |
| AVTU | *Amphora vetula* | Levkov, | 0 | 0 | 0 | 0 |
| AZHA | *Achnanthidium zhakovschikovii* | M. Potapova | 0 | 0 | 0 | 0 |
| BBRE* | *Brachysira brebissonii subsp. brebissonii* | Ross in Hartley | 0 | 0 | 0 | 0 |
| BGAR | *Brachysira garrensis* | (Lange-Bertalot & Krammer) Lange-Bertalot | 0 | 0 | 0 | 0 |
| BLIL | *Brachysira liliana* | Lange-Bertalot | 0 | 0 | 0 | 0 |
| BMIC | *Brachysira microcephala* | (Grunow) Compère | 0 | 0 | 0 | 0 |
| BNEG | *Brachysira* | Lange-Bertalot | 0 | 0 | 0 | 0 |

| code | name | author | Alert taxa (1: yes ; 0: no) BOD5 | SP | NKJ | Pt |
|---|---|---|---|---|---|---|
| | *neglectissima* | | | | | |
| BNEO | *Brachysira neoexilis* | Lange-Bertalot | 0 | 0 | 0 | 0 |
| BPAX | *Bacillaria paxillifera* | (O.F. Müller) Hendey | 0 | 0 | 0 | 0 |
| BPRO | *Brachysira procera* | Lange-Bertalot & Moser | 0 | 0 | 0 | 0 |
| BVIT | *Brachysira vitrea* | (Grunow) Ross in Hartley | 0 | 0 | 0 | 0 |
| CAEX* | *Cymbella excisa var. excisa* | Kützing | 0 | 1 | 0 | 0 |
| CAFF* | *Cymbella affinis var. affinis* | Kützing | 0 | 0 | 0 | 0 |
| CAFM | *Cymbella affiniformis* | Krammer | 0 | 0 | 0 | 0 |
| CAMB | *Craticula ambigua* | (Ehrenberg) Mann | 0 | 0 | 0 | 0 |
| CAPS | *Caloneis alpestris* | (Grunow) Cleve | 0 | 0 | 0 | 0 |
| CATE | *Caloneis tenuis* | (Gregory) Krammer | 0 | 0 | 0 | 0 |
| CATO | *Cyclotella atomus* | Hustedt | 0 | 1 | 1 | 1 |
| CBAC | *Caloneis bacillum* | (Grunow) Cleve | 0 | 0 | 0 | 0 |
| CBAM | *Cymbopleura amphicephala* | Krammer | 0 | 0 | 0 | 0 |
| CBHD | *Cymbopleura hustedtii* | Novelo Tavera & Ibarra | 0 | 0 | 0 | 0 |
| CBKU* | *Cymbopleura kuelbsii var. kuelbsii* | Krammer | 0 | 0 | 0 | 0 |
| CBNA* | *Cymbopleura naviculiformis var. naviculiformis* | (Auerswald) Krammer | 0 | 0 | 0 | 0 |
| CBPY | *Cymbopleura pyrenaica* | Le Cohu et Lange-Bertalot | 0 | 0 | 0 | 0 |
| CCMP | *Cymbella compacta* | Østrup | 0 | 0 | 0 | 0 |
| CCOC | *Cavinula cocconeiformis* | (Gregory ex Greville) Mann & Stickle in Round Crawford & Mann | 0 | 0 | 0 | 0 |
| CCYM | *Cymbella* | Agardh | 0 | 0 | 0 | 0 |

| code | name | author | Alert taxa (1: yes ; O: no) BOD5 | SP | NKJ | Pt |
|---|---|---|---|---|---|---|
| | *cymbiformis* | | | | | |
| CDTG* | *Cyclotella distinguenda var. distinguenda* | Hustedt | 0 | 0 | 0 | 0 |
| CDUB | *Cyclostephanos dubius* | (Fricke) Round | 1 | 1 | 1 | 0 |
| CEUG | *Cocconeis euglypta* | Ehrenberg | 1 | 1 | 1 | 0 |
| CEXF* | *Cymbella excisiformis var. excisiformis* | Krammer | 0 | 0 | 0 | 0 |
| CFDI | *Cymbellafalsa diluviana* | (Krasske) Lange-Bertalot & Metzeltin | 0 | 0 | 0 | 0 |
| CFON | *Caloneis fontinalis* | (Grunow) Lange-Bertalot & Reichardt | 1 | 1 | 1 | 1 |
| CFON | *Caloneis fontinalis* | (Grunow in Van Heurck) Cleve-Euler | 1 | 1 | 1 | 1 |
| CFTF | *Cymbopleura florentiniformis* | Krammer | 0 | 0 | 0 | 0 |
| CHEL | *Cymbella helvetica* | Kützing | 0 | 0 | 0 | 0 |
| CHHA | *Chamaepinnularia hassiaca* | (Krasske) Cantonati & Lange-Bertalot | 0 | 0 | 0 | 0 |
| CHLI | *Craticula halophilioides* | (Hustedt) Lange-Bertalot | 0 | 0 | 0 | 0 |
| CHME | *Chamaepinnularia mediocris* | (Krasske) Lange-Bertalot in Lange-Bertalot & Metzeltin | 0 | 0 | 0 | 0 |
| CINV | *Cyclostephanos invisitatus* | Hohn & Hellerman)Theriot Stoermer & Håkansson | 0 | 0 | 0 | 0 |
| CJAR | *Cavinula jaernefeltii* | (Hustedt) Mann & Stickle in Round Crawford & Mann | 0 | 0 | 0 | 0 |
| CKPP | *Cymbella kappii* | (Cholnoky) Cholnoky | 0 | 0 | 0 | 0 |
| CLAE* | *Cymbella laevis var.* | Naegeli ex Kützing | 0 | 0 | 0 | 0 |

| code | name | author | Alert taxa (1: yes ; O: no) BOD5 | SP | NKJ | Pt |
|---|---|---|---|---|---|---|
| | *laevis* | | | | | |
| CLBE | *Cymbella lange-bertalotii* | Krammer | 0 | 0 | 0 | 0 |
| CLCT | *Caloneis lancettula* | (Schulz) Lange-Bertalot & Witkowski | 1 | 1 | 1 | 1 |
| CLNT | *Cocconeis lineata* | Ehrenberg | 1 | 0 | 0 | 1 |
| CLTL | *Cymbella lancettula* | (Krammer) Krammer | 0 | 0 | 0 | 0 |
| CMDU | *Cyclotella meduanae* | Germain emend Genkal | 1 | 1 | 1 | 1 |
| CMDU | *Cyclotella meduanae* | Germain | 1 | 1 | 1 | 1 |
| CMEN | *Cyclotella meneghiniana* | Kützing | 1 | 1 | 1 | 1 |
| CMLF | *Craticula molestiformis* | (Hustedt) Lange-Bertalot | 0 | 0 | 0 | 0 |
| CNCI* | *Cymbella neocistula var. neocistula* | Krammer | 0 | 0 | 0 | 0 |
| CNLC | *Cymbella neolanceolata* | W. Silva | 0 | 0 | 0 | 0 |
| CNLP* | *Cymbella neoleptoceros var. neoleptoceros* | Krammer | 0 | 1 | 0 | 1 |
| CNTH | *Cocconeis neothumensis* | Krammer | 0 | 0 | 0 | 0 |
| COPL | *Cocconeis pseudolineata* | (Geitler) Lange-Bertalot | 0 | 0 | 0 | 0 |
| CPAR | *Cymbella parva* | (W. Sm.) Kirchner in Cohn | 0 | 0 | 0 | 0 |
| CPED | *Cocconeis pediculus* | Ehrenberg | 0 | 0 | 0 | 0 |
| CPLA* | *Cocconeis placentula var. placentula* | Ehrenberg | 0 | 0 | 0 | 0 |
| CPPV | *Cymbella perparva* | Krammer | 0 | 0 | 0 | 0 |
| CPRX* | *Cymbella proxima var. proxima* | Reimer in Patrick & Reimer | 0 | 0 | 0 | 0 |
| CPSE | *Cavinula pseudoscutiformis* | (Hustedt) Mann & Stickle in Round Crawford | 0 | 0 | 0 | 0 |

| code | name | author | Alert taxa (1: yes ; O: no) | | | |
|---|---|---|---|---|---|---|
| | | | BOD5 | SP | NKJ | Pt |
| | | & Mann | | | | |
| CRAC | *Craticula accomoda* | (Hustedt) Mann | 0 | 0 | 0 | 0 |
| CRBU | *Craticula buderi* | (Hustedt) Lange-Bertalot | 0 | 0 | 0 | 0 |
| CRCU | *Craticula cuspidata* | (Kützing) Mann | 0 | 0 | 0 | 0 |
| CSAQ* | *Cymbopleura subaequalis var. subaequalis* | (Grunow) Krammer | 0 | 0 | 0 | 0 |
| CSBH | *Cymbella subhelvetica* | Krammer | 0 | 0 | 0 | 0 |
| CSCI | *Cymbella subcistula* | Krammer | 0 | 0 | 0 | 0 |
| CSDL | *Cyclostephanos delicatus* | (Genkal) Kling & Håkansson | 0 | 0 | 0 | 0 |
| CSDL | *Cyclostephanos delicatus* | (Genkal) Casper & Scheffler | 0 | 0 | 0 | 0 |
| CSHU | *Caloneis schumanniana* | (Grunow in Van Heurck) Cleve | 0 | 0 | 0 | 0 |
| CSIL | *Caloneis silicula* | (Ehrenberg) Cleve | 0 | 0 | 0 | 0 |
| CSLP | *Cymbella subleptoceros* | Krammer | 1 | 0 | 0 | 1 |
| CSMU | *Chamaepinnularia submuscicola* | (Krasske) Lange-Bertalot | 0 | 0 | 0 | 0 |
| CSNU | *Craticula subminuscula* | (Manguin) C.E. Wetzel & Ector | 1 | 1 | 1 | 1 |
| CSUT* | *Cymbella subtruncata var. subtruncata* | Krammer | 0 | 0 | 0 | 0 |
| CTPU | *Ctenophora pulchella* | (Ralfs ex Kütz.) Williams et Round | 0 | 0 | 0 | 0 |
| CTRQ | *Centric diatoms* | Diatomées centriques indifférenciées | 0 | 0 | 0 | 0 |
| CTUM | *Cymbella tumida* | (Brébisson)Van Heurck | 1 | 1 | 0 | 1 |
| CVMO | *Cavinula mollicula* | (Hustedt) Lange-Bertalot | 0 | 0 | 0 | 0 |
| CVSO | *Cavinula scutelloides* | (W. Smith) Lange-Bertalot | 0 | 0 | 0 | 0 |

| code | name | author | Alert taxa (1: yes ; O: no) BOD5 | SP | NKJ | Pt |
|---|---|---|---|---|---|---|
| CVUL* | *Cymbella vulgata var. vulgata* | Krammer | 0 | 0 | 0 | 0 |
| DCAL | *Diploneis calcilacustris* | Lange-Bertalot et A. Fuhrmann | 0 | 0 | 0 | 0 |
| DCOF* | *Diadesmis confervacea var. confervacea* | Kützing | 0 | 1 | 1 | 1 |
| DEFO* | *Diatomée anormale f. anormale* | Abnormal diatom valve (unidentified) or sum of deformities abundance | 0 | 1 | 1 | 1 |
| DEHR | *Diatoma ehrenbergii* | Kützing | 0 | 0 | 0 | 0 |
| DITE | *Diatoma tenue* | Agardh | 1 | 0 | 0 | 1 |
| DKOT | *Dorofeyukea kotschyi* | (Grunow) Kulikovskiy, Kociolek, Tusset & T.Ludwig | 0 | 0 | 0 | 0 |
| DKRA | *Diploneis krammeri* | Lange-Bertalot & Reichardt | 0 | 0 | 0 | 0 |
| DKUE* | *Denticula kuetzingii var. kuetzingii* | Grunow | 0 | 0 | 0 | 0 |
| DMES | *Diatoma mesodon* | (Ehrenberg) Kützing | 0 | 0 | 0 | 0 |
| DOBL | *Diploneis oblongella* | (Naegeli) Cleve-Euler | 0 | 0 | 0 | 0 |
| DOCU | *Diploneis oculata* | (Brébisson in Desmazières) Cleve | 0 | 0 | 0 | 0 |
| DPAR | *Diploneis parma* | Cleve | 0 | 0 | 0 | 0 |
| DPDE | *Delicatophycus delicatulus* | (Kützing) M.J.Wynne . | 0 | 0 | 0 | 0 |
| DPSG | *Discostella pseudostelligera* | (Hustedt) Houk & Klee emend. Genkal | 0 | 0 | 0 | 1 |
| DPSG | *Discostella pseudostelligera* | (Hustedt) Houk et Klee | 0 | 0 | 0 | 1 |
| DSEP | *Diploneis separanda* | Lange-Bertalot | 0 | 0 | 0 | 0 |
| DSTE | *Discostella* | (Cleve et | 0 | 0 | 0 | 0 |

| code | name | author | Alert taxa (1: yes ; O: no) BOD5 | SP | NKJ | Pt |
|---|---|---|---|---|---|---|
| | *stelligera* | Grun.) Houk & Klee | | | | |
| DTEN | *Denticula tenuis* | Kützing | 0 | 0 | 0 | 0 |
| DVUL | *Diatoma vulgaris* | Bory | 0 | 0 | 0 | 0 |
| EADN | *Epithemia adnata* | (Kützing) Brébisson | 1 | 0 | 1 | 1 |
| EARB | *Eunotia arcubus* | Nörpel-Schempp & Lange-Bertalot | 0 | 0 | 0 | 0 |
| EARC* | *Eunotia arcus var. arcus sensu stricto* | Ehrenberg | 0 | 0 | 0 | 0 |
| EAUE | *Encyonema auerswaldii* | Rabenhorst | 0 | 0 | 0 | 0 |
| EBLU | *Eunotia bilunaris* | (Ehrenberg) M.G.M. Souza in Souza & Moreira-Filho | 0 | 0 | 0 | 0 |
| EBLU | *Eunotia bilunaris* | (Ehrenberg) Schaarschmidt | 0 | 0 | 0 | 0 |
| EBNA | *Encyonema bonapartei* | HeudrE. C.E. Wetzel & Ector | 0 | 0 | 0 | 0 |
| EBOA | *Eunotia boreoalpina* | Lange-Bertalot & Nörpel-Schempp | 0 | 0 | 0 | 0 |
| EBOT | *Eunotia botuliformis* | Wild, Nörpel-Schempp & Lange-Bertalot | 0 | 0 | 0 | 0 |
| EBOT | *Eunotia botuliformis* | Wang | 0 | 0 | 0 | 0 |
| ECAE* | *Encyonema caespitosum var. caespitosum* | Kützing | 0 | 0 | 0 | 0 |
| ECAL | *Encyonopsis alpina* | Krammer & Lange-Bertalot | 0 | 0 | 0 | 0 |
| ECES | *Encyonopsis cesatii* | (Rabenhorst) Krammer | 0 | 0 | 0 | 0 |
| ECKR | *Encyonopsis krammeri* | Reichardt | 0 | 0 | 0 | 0 |
| ECPM | *Encyonopsis minuta* | Krammer & Reichardt | 0 | 0 | 0 | 0 |
| ECTA | *Encyonopsis tavirana* | Krammer | 0 | 0 | 0 | 0 |
| EEXI | *Eunotia exigua* | (Brébisson ex Kützing) Rabenhorst | 0 | 0 | 0 | 0 |

| code | name | author | Alert taxa (1: yes ; O: no) BOD5 | SP | NKJ | Pt |
|---|---|---|---|---|---|---|
| EFAB | *Eunotia faba* | (Ehrenberg) Grunow in Van Heurck | 0 | 0 | 0 | 0 |
| EGBA | *Epithemia gibba* | (Ehrenberg) Kützing | 1 | 0 | 1 | 1 |
| EHOR | *Encyonopsis horticola* | Van de Vijver, Lange-Bertalot & Compère | 0 | 0 | 0 | 0 |
| EIMP | *Eunotia implicata* | Nörpel Lange-Bertalot & Alles | 0 | 0 | 0 | 0 |
| EINC* | *Eunotia incisa var. incisa* | Gregory | 0 | 0 | 0 | 0 |
| ELBV* | *Encyonema lange-bertalotii var. lange-bertalotii* | Krammer | 0 | 0 | 0 | 0 |
| ELEI | *Encyonema leibleinii* | (C. Agardh) Silva, Jahn Ludwig & Menezes | 0 | 1 | 0 | 0 |
| EMIN | *Eunotia minor* | (Kützing) Grunow in Van Heurck | 0 | 0 | 0 | 0 |
| EMIN | *Eunotia minor* | Fusey | 0 | 0 | 0 | 0 |
| EMUC | *Eunotia mucophila* | (Lange-Bert.&Norpel Schempp) Lange-Bertalot | 0 | 0 | 0 | 0 |
| ENAE | *Eunotia naegelii* | Migula | 0 | 0 | 0 | 0 |
| ENCM | *Encyonopsis microcephala* | (Grunow) Krammer | 0 | 0 | 0 | 0 |
| ENEE | *Encyonopsis neerlandica* | Van de Vijver. Verweij, Van Der Wal & Mertens | 0 | 0 | 0 | 0 |
| ENKA | *Encyonema kalbei* | Krammer | 0 | 0 | 0 | 0 |
| ENMI | *Encyonema minutum* | (Hilse in Rabh.) D.G. Mann in Round Crawford & Mann | 0 | 1 | 1 | 1 |
| ENNG | *Encyonema neogracile* | Krammer | 0 | 0 | 0 | 0 |
| ENRE | *Encyonema reichardtii* | (Krammer) D.G. Mann in Round Crawford & Mann | 0 | 0 | 0 | 0 |
| ENRO | *Encyonema rostratum* | Krammer | 0 | 0 | 0 | 0 |

| code | name | author | Alert taxa (1: yes ; 0: no) | | | |
|---|---|---|---|---|---|---|
| | | | BOD5 | SP | NKJ | Pt |
| ENTR | *Encyonema triangulum* | (Ehrenberg) Kützing | 0 | 1 | 0 | 0 |
| ENVE | *Encyonema ventricosum* | (Kützing) Grunow in Schmidt & al. | 0 | 0 | 0 | 0 |
| EOCO | *Eolimna comperei* | Ector Coste et Iserentant in Coste & Ector | 0 | 0 | 0 | 0 |
| EORT | *Eunotia orthohedra* | Furey, Lowe et Johansen | 0 | 0 | 0 | 0 |
| EPBO | *Epithemia proboscidea* | Kützing | 0 | 0 | 0 | 0 |
| EPEC* | *Eunotia pectinalis var. pectinalis* | (Kützing) Rabenhorst | 0 | 0 | 0 | 0 |
| EPHP | *Epithemia parallela* | (Grunow) Ruck & Nakov | 0 | 0 | 0 | 0 |
| EPHP | *Epithemia parallela* | Proschkina-Lavrenko | 0 | 0 | 0 | 0 |
| EREI | *Epithemia reicheltii* | Fricke | 0 | 0 | 0 | 0 |
| ERHO | *Eunotia rhomboidea* | Hustedt | 0 | 0 | 0 | 0 |
| ESLE | *Encyonema silesiacum* | (Bleisch in Rabh.) D.G. Mann | 0 | 1 | 0 | 1 |
| ESMI | *Epithemia smithii* | Carruthers in Gray | 0 | 0 | 0 | 0 |
| ESOR | *Epithemia sorex* | Kützing | 1 | 1 | 1 | 1 |
| ESUB | *Eunotia subarcuatoides* | Alles Nörpel & Lange-Bertalot in Alles et al. | 0 | 0 | 0 | 0 |
| ESUM | *Encyonopsis subminuta* | Krammer & Reichardt | 0 | 0 | 0 | 0 |
| ETEN | *Eunotia tenella* | (Grunow in Van Heurck) Hustedt in Schmidt & al | 0 | 0 | 0 | 0 |
| ETUR* | *Epithemia turgida var. turgida* | (Ehrenberg) Kützing | 0 | 0 | 0 | 0 |
| EUAL | *Eucocconeis alpestris* | (Brun) Lange-Bertalot | 0 | 0 | 0 | 0 |
| EUBI | *Eunotia bidens* | Ehrenberg | 0 | 0 | 0 | 0 |
| EUFL | *Eucocconeis flexella* | (Kützing) Meister | 0 | 0 | 0 | 0 |
| EULA | *Eucocconeis* | (Østrup) | 0 | 0 | 0 | 0 |

| code | name | author | Alert taxa (1: yes ; O: no) BOD5 | SP | NKJ | Pt |
|---|---|---|---|---|---|---|
| | *laevis* | Lange-Bertalot | | | | |
| EVUL* | *Encyonema vulgare var. vulgare* | Krammer | 0 | 1 | 0 | 0 |
| FAPO | *Fragilaria amphicephaloides* | Lange-Bertalot in Hofmann & al. | 0 | 0 | 0 | 0 |
| FAQU | *Fragilaria aquaplus* | Lange-Bertalot & Ulrich | 0 | 0 | 0 | 0 |
| FAUT | *Fragilaria austriaca* | (Grunow) Lange-Bertalot | 0 | 0 | 0 | 0 |
| FCRO | *Fragilaria crotonensis* | Kitton | 0 | 0 | 0 | 0 |
| FCRS | *Frustulia crassinervia* | (Breb.) Lange-Bertalot et Krammer | 0 | 0 | 0 | 0 |
| FERI | *Frustulia erifuga* | Lange-Bertalot & Krammer | 0 | 0 | 0 | 0 |
| FFBI | *Fragilariforma bicapitata* | (A.Mayer) Williams & Round | 0 | 0 | 0 | 0 |
| FFNI | *Fragilariforma nitzschioides* | (Grunow) Lange-Bertalot in Hofmann Werum & Lange-Bertalot | 0 | 0 | 0 | 0 |
| FFUN | *Fragilariforma undata* | (W.Smith) Heudre, C.E.Wetzel & Ector | 0 | 0 | 0 | 0 |
| FFUS | *Fragilaria fusa* | (R.M. Patrick) Wengrat, C.E. Wetzel & E. Morales | 1 | 1 | 0 | 1 |
| FFVI | *Fragilariforma virescens* | (Ralfs) Williams & Round | 0 | 0 | 0 | 0 |
| FGRA | *Fragilaria gracilis* | Østrup | 0 | 0 | 0 | 0 |
| FLEN | *Fallacia lenzii* | Hustedt) Lange-Bertalot | 0 | 0 | 0 | 0 |
| FLEN | *Fallacia lenzii* | (Hustedt) Mann in Van de Vijver & al | 0 | 0 | 0 | 0 |
| FMES | *Fragilaria mesolepta* | Rabenhorst | 0 | 0 | 0 | 0 |
| FMIT | *Fallacia mitis* | (Hustedt) D.G. Mann | 0 | 0 | 0 | 0 |
| FMIV | *Fragilaria microvaucheri* | C.E. Wetzel et Ector | 0 | 0 | 0 | 0 |

| code | name | author | Alert taxa (1: yes ; 0: no) BOD5 | SP | NKJ | Pt |
|---|---|---|---|---|---|---|
| | *ae* | | | | | |
| FNEV | *Fragilaria nevadensis* | Linares-Cuesta & Sanchez-Castillo | 0 | 0 | 0 | 0 |
| FNIN | *Fragilaria neointermedia* | Tuji et D.M. Williams | 0 | 0 | 0 | 0 |
| FPDE | *Fragilaria perdelicatissima* | Lange-Bertalot & Van de Vijver | 0 | 0 | 0 | 0 |
| FPEC | *Fragilaria pectinalis* | Lyngbye | 0 | 1 | 0 | 1 |
| FPEM | *Fragilaria perminuta* | (Grunow) Lange-Bertalot | 0 | 0 | 0 | 0 |
| FPRU | *Fragilaria pararumpens* | Lange-Bertalot, Hofmann & Werum in Hofmann & al. | 0 | 1 | 1 | 1 |
| FRAD | *Fragilaria radians* | (Kütz.) Williams & Round | 0 | 0 | 0 | 0 |
| FRAD | *Fragilaria radians* | Lange-Bertalot in Hofmann & al. | 0 | 0 | 0 | 0 |
| FRUM | *Fragilaria rumpens* | (Kütz.) G.W.F. Carlson | 0 | 1 | 0 | 1 |
| FSAP | *Fistulifera saprophila* | (Lange-Bertalot & Bonik) Lange-Bertalot | 0 | 0 | 0 | 0 |
| FSAX | *Frustulia saxonica* | Rabenhorst | 0 | 0 | 0 | 0 |
| FSBH | *Fallacia subhamulata* | (Grunow in V. Heurck) D.G. Mann | 0 | 0 | 0 | 0 |
| FSCS | *Fragilaria subconstricta* | Østrup | 0 | 0 | 0 | 0 |
| FSCS | *Fragilaria subconstricta* | Østrup (Oestrup 1910) emend Heudre | 0 | 0 | 0 | 0 |
| FSLU | *Fallacia sublucidula* | (Hustedt) D.G. Mann | 0 | 0 | 0 | 0 |
| FSOC | *Fragilaria socia* | (Wallace) Lange-Bertalot | 0 | 0 | 0 | 0 |
| FSXP | *Fragilaria saxoplanctonica* | Lange-Bertalot & Ulrich | 0 | 0 | 0 | 0 |
| FTEN | *Fragilaria tenera* | (W. Smith) Lange-Bertalot | 0 | 0 | 0 | 0 |
| FTNU | *Fragilaria tenuissima* | Lange-Bertalot & Ulrich | 0 | 0 | 0 | 0 |

| | | | Alert taxa (1: yes ; O: no) | | | |
|---|---|---|---|---|---|---|
| code | name | author | BOD5 | SP | NKJ | Pt |
| FVAU* | *Fragilaria vaucheriae var. vaucheriae* | (Kützing) Petersen | 1 | 1 | 1 | 1 |
| FVUL | *Frustulia vulgaris* | (Thwaites) De Toni | 0 | 0 | 0 | 0 |
| GACC | *Geissleria acceptata* | (Hustedt) Lange-Bertalot & Metzeltin | 0 | 0 | 0 | 0 |
| GACD | *Gomphonema acidoclinatiforme* | Metzeltin & Lange-Bertalot | 0 | 0 | 0 | 0 |
| GACU* | *Gomphonema acuminatum var. acuminatum* | Ehrenberg | 0 | 0 | 0 | 0 |
| GADC | *Gomphonema acidoclinatum* | Lange-Bertalot & Reichardt | 1 | 1 | 1 | 1 |
| GAFF | *Gomphonema affine* | Kützing | 0 | 0 | 0 | 0 |
| GAGU | *Gomphonema angustius* | E. Reichardt | 0 | 0 | 0 | 0 |
| GAGV | *Gomphonema angustivalva* | E. Reichardt | 0 | 0 | 0 | 0 |
| GANG | *Gomphonema angustatum* | (Kützing) Rabenhorst | 0 | 0 | 1 | 1 |
| GANT | *Gomphonema angustum* | Agardh sensu Reichardt & Lange Bertalot | 0 | 0 | 0 | 0 |
| GANT | *Gomphonema angustum* | Agardh | 0 | 0 | 0 | 0 |
| GAUG | *Gomphonema augur* | Ehrenberg | 0 | 0 | 0 | 0 |
| GAUR | *Gomphonema auritum* | A. Braun ex Kützing | 0 | 0 | 0 | 0 |
| GBOB | *Gomphonema bourbonense* | E. Reichardt et Lange-Bertalot | 1 | 1 | 1 | 1 |
| GBRE | *Gomphonema brebissonii* | Kützing | 0 | 0 | 0 | 0 |
| GCAD | *Gomphonema campodunense* | E.Reichardt | 0 | 0 | 1 | 0 |
| GCAP | *Gomphonema capitatum* | Ehrenberg | 0 | 0 | 0 | 0 |
| GCLA | *Gomphonema clavatum* | Ehrenberg | 0 | 0 | 0 | 1 |
| GCOR | *Gomphonema coronatum* | Ehrenberg | 0 | 0 | 0 | 0 |
| GCUN | *Gomphonema cuneolus* | E. Reichardt | 0 | 0 | 0 | 0 |

| | | | Alert taxa (1: yes ; O: no) | | | |
|---|---|---|---|---|---|---|
| code | name | author | BOD5 | SP | NKJ | Pt |
| GCUV | *Gomphonema curvipedatum* | H. Kobayasi ex Osada | 0 | 0 | 0 | 0 |
| GCUW | *Geissleria cummerowi* | (L. Kalbe) Lange-Bertalot | 0 | 0 | 0 | 0 |
| GELG | *Gomphonema elegantissimum* | Reichardt & Lange-Bertalot in Hofmann & al. | 0 | 0 | 0 | 0 |
| GERI | *Gomphoneis eriense* | (Grunow) Skvortzow & Meyer | 0 | 0 | 0 | 0 |
| GEXL | *Gomphonema exilissimum* | (Grun.) Lange-Bertalot & Reichardt | 0 | 0 | 0 | 0 |
| GGDI | *Gomphonema graciledictum* | E.Reichardt | 0 | 0 | 1 | 0 |
| GGRA | *Gomphonema gracile* | Ehrenberg | 0 | 0 | 0 | 0 |
| GHEB | *Gomphonema hebridense* | Gregory | 0 | 0 | 0 | 0 |
| GITA | *Gomphonema italicum* | Kützing | 0 | 0 | 0 | 0 |
| GLAT | *Gomphonema lateripunctatum* | Reichardt & Lange-Bertalot | 0 | 0 | 0 | 0 |
| GLGN | *Gomphonema lagenula* | Kützing | 0 | 0 | 0 | 0 |
| GLOV | *Gomphonella olivacea* | NA | 1 | 1 | 0 | 0 |
| GLTC | *Gomphonema laticollum* | Reichardt | 0 | 0 | 0 | 0 |
| GMEX | *Gomphonema mexicanum* | Grunow | 0 | 0 | 0 | 0 |
| GMIC* | *Gomphonema micropus var. micropus* | Kützing | 0 | 0 | 0 | 0 |
| GMIN* | *Gomphonema minutum f. minutum* | (Ag.)Agardh | 1 | 1 | 1 | 1 |
| GMIS | *Gomphonema minusculum* | Krasske | 0 | 0 | 0 | 0 |
| GNLC | *Gomphonella calcarea* | (Cleve) R.Jahn & N.Abarca, comb. nov. | 0 | 0 | 0 | 0 |
| GNVC | *Gomphonema naviculoides* | W. Smith | 0 | 0 | 0 | 0 |
| GOCU | *Gomphonema occultum* | Reichardt & Lange-Bertalot | 0 | 0 | 0 | 0 |
| GOLD | *Gomphonema* | Hustedt | 0 | 0 | 0 | 0 |

| code | name | author | Alert taxa (1: yes ; O: no) BOD5 | SP | NKJ | Pt |
|---|---|---|---|---|---|---|
| | *olivaceoides* | | | | | |
| GPAN | *Gomphocymbellopsis ancyli* | (Cleve) Krammer | 0 | 0 | 0 | 0 |
| GPAR* | *Gomphonema parvulum var. parvulum f. parvulum* | (Kützing) Kützing | 1 | 1 | 1 | 1 |
| GPLI | *Gomphosphenia lingulatiformis* | (Lange-Bertalot & Reichardt) Lange-Bertalot | 0 | 1 | 0 | 0 |
| GPSA | *Gomphonema pseudoaugur* | Lange-Bertalot | 0 | 0 | 0 | 0 |
| GPUM | *Gomphonema pumilum* | (Grunow) Reichardt & Lange-Bertalot | 1 | 1 | 1 | 1 |
| GRHB | *Gomphonema rhombicum* | M. Schmidt | 0 | 0 | 0 | 0 |
| GRHB | *Gomphonema rhombicum* | Fricke | 0 | 0 | 0 | 0 |
| GSBG | *Gomphonema subangustatum* | Lange-Bertalot Cavacini Tagliaventi & Alfinito | 0 | 0 | 0 | 0 |
| GSCI | *Gyrosigma sciotoense* | (Sullivan et Wormley) Cleve | 0 | 0 | 0 | 0 |
| GSCL | *Gomphonema subclavatum* | Grunow | 1 | 1 | 1 | 1 |
| GSPP | *Gomphonema saprophilum* | (Lange-Bertalot & Reichardt) Abarca, R. Jahn, J. Zimmermann & Enke | 0 | 1 | 1 | 1 |
| GTER | *Gomphonema tergestinum* | (Grunow in Van Heurck) Schmidt in Schmidt & al. | 0 | 0 | 0 | 0 |
| GTNO | *Gomphonema tenoccultum* | Reichardt | 0 | 0 | 0 | 0 |
| GTRU | *Gomphonema truncatum* | Ehrenberg | 0 | 0 | 0 | 0 |
| GVIB | *Gomphonema vibrio* | Ehrenberg | 0 | 0 | 0 | 0 |
| GVRD | *Gomphonema varioreduncum* | Jüttner, Ector, Reichardt, Van de Vijver & Cox | 0 | 0 | 0 | 0 |
| GYAT | *Gyrosigma attenuatum* | (Kützing) Rabenhorst | 0 | 0 | 0 | 0 |
| GYKU | *Gyrosigma* | (Grunow) Cleve | 0 | 0 | 0 | 0 |

| code | name | author | Alert taxa (1: yes ; O: no) | | | |
|---|---|---|---|---|---|---|
| | | | BOD5 | SP | NKJ | Pt |
| | *kuetzingii* | | | | | |
| HARC | *Hannaea arcus* | (Ehr.)Patrick | 0 | 0 | 0 | 0 |
| HCAP | *Hippodonta capitata* | (Ehr.)Lange-Bert.Metzeltin & Witkowski | 0 | 1 | 0 | 0 |
| HCOS | *Hippodonta costulata* | (Grunow)Lange-Bertalot Metzeltin & Witkowski | 0 | 1 | 0 | 1 |
| HLMO | *Halamphora montana* | (Krasske) Levkov, | 0 | 0 | 0 | 0 |
| HNEG | *Hippodonta neglecta* | Lange-Bertalot Metzeltin & Witkowski | 0 | 0 | 0 | 0 |
| HOLI | *Halamphora oligotraphenta* | (Lange-Bertalot) Levkov | 0 | 0 | 0 | 0 |
| HPDA | *Hippodonta pseudacceptata* | (Kobayasi) Lange-Bertalot Metzeltin & Witkowski | 0 | 0 | 0 | 0 |
| HPEP | *Humidophila perpusilla* | (Grunow) Lowe, Kociolek, Johansen,Van de Vijver, Lange-Bertalot & Kopalová | 0 | 0 | 0 | 0 |
| HSMA | *Humidophila schmassmannii* | (Hustedt) Buczkó et Wojtal | 0 | 0 | 0 | 0 |
| HTHU | *Halamphora thumensis* | (A.Mayer) Levkov | 0 | 0 | 0 | 0 |
| HUCO | *Humidophila contenta* | (Grunow) Lowe, Kociolek, Johansen, Van de Vijver, Lange-Bertalot & Kopalová | 0 | 0 | 0 | 0 |
| HVEN | *Halamphora veneta* | (Kützing) Levkov, | 0 | 0 | 0 | 0 |
| IDEL | *Iconella delicatissima* | (Lewis) Ruck & Nakov | 0 | 0 | 0 | 0 |
| KALA | *Karayevia laterostrata* | (Hustedt) Bukhtiyarova | 0 | 0 | 0 | 0 |
| KALA | *Karayevia laterostrata* | (Hustedt) Kingston | 0 | 0 | 0 | 0 |
| KAPL | *Karayevia ploenensis* | (Hustedt) Bukhtiyarova | 0 | 0 | 0 | 0 |
| KCLE* | *Karayevia clevei var.* | (Grunow) Bukhtiyarova | 0 | 0 | 0 | 0 |

| code | name | author | Alert taxa (1: yes ; O: no) | | | |
|---|---|---|---|---|---|---|
| | | | BOD5 | SP | NKJ | Pt |
| | *clevei* | | | | | |
| LBAL | *Lindavia balatonis* | (Pantocsek) Nakov, Guillory, Julius, Theriot & Alverson | 0 | 0 | 0 | 0 |
| LGOP | *Luticola goeppertiana* | (Bleisch in Rabenhorst)D.G. Mann in Round Crawford & Mann | 0 | 0 | 0 | 0 |
| LGOP | *Luticola goeppertiana* | (Bleisch) D.G.Mann ex J.Rarick, S.Wu, S.S.Lee & Edlund | 0 | 0 | 0 | 0 |
| LHUN | *Lemnicola hungarica* | (Grunow) Round & Basson | 0 | 0 | 0 | 0 |
| LPRA | *Lindavia praetermissa* | (Lund) Nakov, Guillory, Julius, Theriot & Alverson | 0 | 0 | 0 | 0 |
| LRAD | *Lindavia radiosa* | (Grunow) De Toni & Forti | 0 | 0 | 0 | 0 |
| MAAT* | *Mayamaea atomus var. atomus* | (Kützing) Lange-Bertalot | 0 | 0 | 0 | 0 |
| MALC | *Mayamaea alcimonica* | (E. Reichardt) C.E. Wetzel, Barragán & Ector | 0 | 0 | 0 | 0 |
| MCIR* | *Meridion circulare var. circulare* | (Greville) C.A. Agardh | 0 | 0 | 0 | 0 |
| MING | *Mayamaea ingenua* | (Hustedt) Lange-Bertalot & Hofmann in Hofmann & al. | 0 | 1 | 1 | 0 |
| MLAC | *Mastogloia lacustris* | (Grunow) van Heurck | 0 | 0 | 0 | 0 |
| MPMI | *Mayamaea permitis* | (Hustedt) Bruder & Medlin | 0 | 1 | 1 | 1 |
| MSMI | *Mastogloia smithii* | Thwaites | 0 | 0 | 0 | 0 |
| MSTJ | *Mastogloia sterijovskii* | A. Pavlov. Jovanovska, C.E.Wetzel, Ector & Levkov | 0 | 0 | 0 | 0 |
| MVAR | *Melosira varians* | Agardh | 1 | 1 | 1 | 1 |

| code | name | author | Alert taxa (1: yes ; O: no) | | | |
|---|---|---|---|---|---|---|
| | | | BOD5 | SP | NKJ | Pt |
| NAAN | *Navicula angusta* | Grunow | 0 | 0 | 0 | 0 |
| NACD | *Nitzschia acidoclinata* | Lange-Bertalot | 0 | 0 | 0 | 0 |
| NACI | *Nitzschia acicularis* | Kützing) W.M.Smith | 1 | 1 | 1 | 1 |
| NACU | *Nitzschia acula* | Hantzsch ex Cleve & Grunow | 0 | 0 | 0 | 0 |
| NAGN | *Nitzschia agnita* | Hustedt | 0 | 0 | 0 | 0 |
| NAGW | *Nitzschia agnewii* | Cholnoky | 0 | 0 | 0 | 0 |
| NALP | *Neidium alpinum* | Hustedt | 0 | 0 | 0 | 0 |
| NAMP* | *Nitzschia amphibia f. amphibia* | Grunow | 1 | 1 | 1 | 1 |
| NANT | *Navicula antonii* | Lange-Bertalot | 1 | 1 | 1 | 1 |
| NAPB | *Nitzschia alpinobacillum* | Lange-Bertalot | 0 | 0 | 0 | 0 |
| NCAR | *Navicula cari* | Ehrenberg | 0 | 0 | 0 | 0 |
| NCAT | *Navicula catalanogermanica* | Lange-Bertalot & Hofmann | 0 | 0 | 0 | 0 |
| NCIN | *Navicula cincta* | (Ehr.) Ralfs in Pritchard | 0 | 0 | 0 | 0 |
| NCLA | *Nitzschia clausii* | Hantzsch | 0 | 0 | 0 | 0 |
| NCPL | *Nitzschia capitellata* | Hustedt in A. Schmidt & al. | 1 | 1 | 1 | 1 |
| NCPR | *Navicula capitatoradiata* | Germain | 1 | 1 | 1 | 1 |
| NCRY | *Navicula cryptocephala* | Kützing | 0 | 0 | 1 | 0 |
| NCTE | *Navicula cryptotenella* | Lange-Bertalot | 0 | 0 | 0 | 0 |
| NCTO | *Navicula cryptotenelloides* | Lange-Bertalot | 0 | 1 | 0 | 0 |
| NCTT | *Navicula cataracta-rheni* | Lange-Bertalot | 1 | 1 | 0 | 1 |
| NCTV | *Navicula caterva* | Hohn & Hellerman | 0 | 1 | 0 | 0 |
| NDBF | *Neidiomorpha binodeformis* | Cantonati, Lange-Bertalot | 0 | 0 | 0 | 0 |

| code | name | author | Alert taxa (1: yes ; 0: no) | | | |
|---|---|---|---|---|---|---|
| | | | BOD5 | SP | NKJ | Pt |
| | | & Angeli | | | | |
| NDIS* | *Nitzschia dissipata subsp. dissipata* | (Kützing) Grunow | 0 | 1 | 0 | 0 |
| NDRA | *Nitzschia draveillensis* | Coste & Ricard | 0 | 1 | 1 | 1 |
| NEDU | *Neidium dubium* | (Ehrenberg)Cleve | 0 | 0 | 0 | 0 |
| NERI | *Navicula erifuga* | Lange-Bertalot in Krammer & Lange-Bertalot | 0 | 0 | 0 | 0 |
| NEUT | *Nitzschia eutinensis* | Lange-Bertalot & Werum | 0 | 0 | 0 | 0 |
| NEXI | *Navicula exilis* | Kützing | 0 | 0 | 0 | 0 |
| NFIL* | *Nitzschia filiformis var. filiformis* | (W.M.Smith) Van Heurck | 1 | 1 | 0 | 1 |
| NFON | *Nitzschia fonticola* | Grunow in Cleve et Möller | 0 | 0 | 0 | 0 |
| NFSO | *Nanofrustulum sopotensis* | (Witkowski & Lange-Bert.) E.Morales, C.E.Wetzel & Ector, comb. nov. | 0 | 0 | 0 | 1 |
| NFTR | *Nanofrustulum trainori* | (E.Morales) E.Morales, comb. nov. | 1 | 1 | 0 | 1 |
| NGDU | *Navigeia decussis* | (Østrup) Bukhtiyarova | 0 | 0 | 0 | 0 |
| NGER | *Navicula germainii* | Wallace | 0 | 1 | 0 | 1 |
| NGES | *Nitzschia gessneri* | Hustedt | 0 | 0 | 0 | 0 |
| NGHI | *Navigeia hinziae* | (Novais et Ector) Bukhtiyarova | 0 | 0 | 0 | 0 |
| NGOT | *Navicula gottlandica* | Grunow in Van Heurck | 0 | 0 | 0 | 0 |
| NGRE | *Navicula gregaria* | Donkin | 0 | 0 | 0 | 1 |
| NHAN | *Nitzschia hantzschiana* | Rabenhorst | 0 | 0 | 0 | 0 |
| NHEU | *Nitzschia heufleriana* | Grunow | 0 | 0 | 0 | 0 |
| NHIN | *Navicula hintzii* | Lange-Bertalot | 0 | 1 | 0 | 1 |

| | | | Alert taxa (1: yes ; 0: no) | | | |
|---|---|---|---|---|---|---|
| code | name | author | BOD5 | SP | NKJ | Pt |
| NHMD | *Navicula heimansioides* | Lange-Bertalot | 0 | 0 | 0 | 0 |
| NIAR | *Nitzschia archibaldii* | Lange-Bertalot | 0 | 0 | 0 | 0 |
| NIBU | *Nitzschia bulnheimiana* | (Rabenhorst) H.L.Smith | 0 | 0 | 0 | 0 |
| NIFQ | *Nitzschia frequens* | Hustedt | 0 | 1 | 0 | 1 |
| NIFR* | *Nitzschia frustulum var. frustulum* | (Kützing) Grunow | 1 | 1 | 1 | 1 |
| NIFT | *Nitzschia fruticosa* | Hustedt | 0 | 0 | 0 | 0 |
| NIGR | *Nitzschia gracilis* | Hantzsch | 1 | 1 | 1 | 1 |
| NILA | *Nitzschia lacuum* | Lange-Bertalot | 0 | 0 | 0 | 0 |
| NIME | *Nitzschia media* | Hantzsch. | 0 | 0 | 0 | 0 |
| NINC | *Nitzschia inconspicua* | Grunow | 1 | 1 | 1 | 1 |
| NINT | *Nitzschia intermedia* | Hantzsch ex Cleve & Grunow | 1 | 1 | 1 | 1 |
| NIOG | *Nitzschia oligotraphenta* | (Lange-Bertalot) Lange-Bertalot in Hofmann & al. | 0 | 0 | 0 | 0 |
| NIPF | *Nitzschia paleaeformis* | Hustedt | 0 | 0 | 0 | 0 |
| NIPM | *Nitzschia perminuta* | (Grunow) M.Peragallo | 0 | 0 | 0 | 0 |
| NISO | *Nitzschia solita* | Hustedt | 0 | 0 | 0 | 0 |
| NISU | *Nitzschia subtilis* | Grunow in Cleve et Grunow | 0 | 0 | 0 | 0 |
| NIVA | *Nitzschia valdestriata* | Aleem & Hustedt | 0 | 0 | 0 | 0 |
| NJOC | *Navicula johncarterii* | D.M.Williams | 0 | 0 | 0 | 0 |
| NLAL | *Nitzschia labella* | Moser Lange-Bertalot & Metzeltin | 0 | 0 | 0 | 0 |
| NLAN | *Navicula lanceolata* | (Agardh) Ehrenberg | 0 | 0 | 0 | 0 |
| NLAN | *Navicula lanceolata* | (Agardh) Kützing | 0 | 0 | 0 | 0 |
| NLIN* | *Nitzschia linearis var.* | (Agardh) W.M.Smith | 0 | 0 | 0 | 0 |

| code | name | author | Alert taxa (1: yes ; 0: no) | | | |
|---|---|---|---|---|---|---|
| | | | BOD5 | SP | NKJ | Pt |
| | *linearis* | | | | | |
| NLTK | *Navicula leistikowii* | Lange-Bertalot | 0 | 0 | 0 | 0 |
| NLUN | *Navicula lundii* | Reichardt | 0 | 0 | 0 | 0 |
| NMCA | *Navicula microcari* | Lange-Bertalot | 0 | 0 | 0 | 0 |
| NMEN* | *Navicula menisculus var. menisculus* | Schumann | 0 | 0 | 0 | 0 |
| NMIC | *Nitzschia microcephala* | Grunow in Cleve & Moller | 0 | 0 | 0 | 0 |
| NMOK | *Navicula moskalii* | Metzeltin, Witkowski & Lange-Bertalot | 0 | 0 | 0 | 0 |
| NMTA | *Navicula metareichardtiana* | Lange-Bertalot & Kusber nom.nov. | 0 | 1 | 0 | 1 |
| NNAN | *Nitzschia nana* | Grunow in Van Heurck | 0 | 0 | 0 | 0 |
| NNOT | *Navicula notha* | Wallace | 0 | 0 | 0 | 0 |
| NOBL | *Navicula oblonga* | Kützing | 0 | 0 | 0 | 0 |
| NOLI | *Navicula oligotraphenta* | Lange-Bertalot & Hofmann | 0 | 0 | 0 | 0 |
| NPAE | *Nitzschia paleacea* | (Grunow) Grunow in Van Heurck | 1 | 1 | 1 | 1 |
| NPAL* | *Nitzschia palea var. palea* | (Kützing) W.Smith | 1 | 1 | 0 | 1 |
| NPML | *Nitzschia pumila* | Hustedt | 1 | 1 | 1 | 1 |
| NPRA | *Navicula praeterita* | Hustedt | 0 | 0 | 0 | 0 |
| NPSL | *Navicula pseudolanceolata* | Lange-Bertalot | 0 | 0 | 0 | 0 |
| NPUF | *Nitzschia puriformis* | Hlubikova et Ector | 0 | 0 | 0 | 0 |
| NRAD | *Navicula radiosa* | Kützing | 0 | 0 | 0 | 0 |
| NRCS | *Navicula recens* | (Lange-Bertalot) Lange-Bertalot | 0 | 0 | 0 | 0 |
| NREC | *Nitzschia recta* | Hantzsch in Rabenhorst | 0 | 0 | 0 | 0 |

| code | name | author | Alert taxa (1: yes ; O: no) | | | |
|---|---|---|---|---|---|---|
| | | | BOD5 | SP | NKJ | Pt |
| NRFA | *Navicula radiosafallax* | Lange-Bertalot | 0 | 0 | 0 | 0 |
| NRHT | *Navicula rhynchotella* | Lange-Bertalot | 0 | 0 | 0 | 0 |
| NRHY | *Navicula rhynchocephala* | Kützing | 0 | 0 | 0 | 0 |
| NROS | *Navicula rostellata* | Kützing | 0 | 0 | 0 | 0 |
| NSBL | *Nitzschia sublinearis* | Hustedt | 0 | 0 | 0 | 0 |
| NSBN | *Navicula subalpina* | Reichardt | 0 | 0 | 0 | 0 |
| NSIA | *Navicula simulata* | Manguin | 1 | 1 | 1 | 1 |
| NSNM | *Navicula sancti-naumii* | Levkov, et Metzeltin | 0 | 0 | 0 | 0 |
| NSOC | *Nitzschia sociabilis* | Hustedt | 0 | 1 | 0 | 0 |
| NSOL | *Nitzschia solgensis* | Cleve-Euler | 1 | 1 | 1 | 1 |
| NSTS | *Nitzschia soratensis* | Morales & Vis | 1 | 1 | 0 | 1 |
| NSUA | *Nitzschia subacicularis* | Hustedt in A. Schmidt et al. | 1 | 1 | 1 | 1 |
| NTAB | *Nitzschia tabellaria* | (Grun.) Grun. in Cl. & Grunow | 0 | 1 | 0 | 0 |
| NTCX | *Navicula trophicatrix* | Lange-Bertalot | 0 | 0 | 0 | 0 |
| NTPT | *Navicula tripunctata* | (O.F.Müller) Bory | 1 | 1 | 1 | 1 |
| NTRV* | *Navicula trivialis var. trivialis* | Lange-Bertalot | 0 | 0 | 0 | 0 |
| NUSA | *Navicula upsaliensis* | (Grunow) Peragallo | 0 | 0 | 0 | 0 |
| NUVI | *Nupela vitiosa* | (Schimanski) Lange-Bertalot in Krammer & Lange-Bertalot | 0 | 0 | 0 | 0 |
| NUWE | *Nupela wellneri* | (Lange-Bertalot) Lange-Bertalot | 0 | 0 | 0 | 0 |
| NVDA* | *Navicula vandamii var. vandamii* | Schoeman & Archibald | 0 | 1 | 0 | 0 |
| NVEN | *Navicula veneta* | Kützing | 0 | 1 | 1 | 1 |
| NVGL | *Navicula* | Hustedt | 0 | 0 | 0 | 0 |

| | | | Alert taxa (1: yes ; O: no) | | | |
|---|---|---|---|---|---|---|
| code | name | author | BOD5 | SP | NKJ | Pt |
| | *virginalis* | | | | | |
| NVIR* | *Navicula viridula var. viridula* | (Kützing) Ehrenberg | 0 | 0 | 0 | 0 |
| NVXP | *Nitzschia vixpalea* | Lange-Bertalot & Werum | 0 | 0 | 0 | 0 |
| NWIL | *Navicula wildii* | Lange-Bertalot | 0 | 0 | 0 | 0 |
| NXAS | *Navicula associata* | Lange-Bertalot | 0 | 0 | 0 | 0 |
| NYCO | *Nitzschia costei* | Tudesque, Rimet & Ector | 0 | 0 | 0 | 0 |
| NZAL | *Nitzschia alpina* | Hustedt | 0 | 0 | 0 | 0 |
| NZRA | *Nitzschia radicula* | Hustedt | 0 | 0 | 0 | 0 |
| NZSU | *Nitzschia supralitorea* | Lange-Bertalot | 0 | 1 | 0 | 0 |
| PABV | *Planothidium abbreviatum* | (Reimer) Potapova | 1 | 0 | 0 | 1 |
| PADE | *Pantocsekiell a delicatula* | (Hustedt) K.T. Kiss et Ács | 0 | 0 | 0 | 0 |
| PALT | *Psammothidium altaicum* | (Poretzky) Bukhtiyarova in Bukhtiyarova & Round | 0 | 0 | 0 | 0 |
| PALV | *Pseudostauros ira alvareziae* | Cejudo-Figuera s Morales & Ector | 0 | 0 | 0 | 1 |
| PCIT | *Playaensis citrus* | (Krasske) E. Reichardt nov. comb. | 0 | 0 | 0 | 0 |
| PCLD | *Placoneis clementioides* | (Hustedt) Cox | 0 | 0 | 0 | 0 |
| PCLT | *Placoneis clementis* | (Grunow) Cox | 0 | 0 | 0 | 0 |
| PCMS | *Pantocsekiell a comensis* | (Grunow in Van Heurck) K.T. Kiss et Ács | 0 | 0 | 0 | 0 |
| PCOS | *Pantocsekiell a costei* | (Druart et F. Straub) K.T. Kiss et Ács | 0 | 0 | 0 | 0 |
| PDAU | *Planothidium daui* | (Foged) Lange-Bertalot | 0 | 1 | 0 | 0 |
| PDID | *Psammothidium didymum* | (Hustedt ) Bukhtiyarova et Round | 0 | 0 | 0 | 0 |
| PDOP | *Pseudostauros* | (Lange-Bertalo | 0 | 0 | 0 | 0 |

| code | name | author | Alert taxa (1: yes ; O: no) BOD5 | SP | NKJ | Pt |
|---|---|---|---|---|---|---|
| | *ira parasitoides* | t, Rol.Schmidt & Klee in Schmidt et al.) E.Morales, M.L.García & Maidana | | | | |
| PDPC | *Pseudostauros iropsis connecticuten sis* | Morales | 1 | 0 | 0 | 1 |
| PENA | *Pennate diatom diatomée pennée non identifiée* | in Kelly application TDI | 0 | 0 | 0 | 0 |
| PFIB | *Peronia fibula* | (Brébisson ex Kützing)Ross | 0 | 0 | 0 | 0 |
| PGRI | *Psammothidium grischunum* | (Wuthrich) Bukhtiyarova et Round | 0 | 0 | 0 | 0 |
| PGRN | *Planothidium granum* | (Hohn & Hellerman) Lange-Bertalot | 1 | 0 | 0 | 1 |
| PHEL | *Psammothidium helveticum* | (Hustedt) Bukhtiyarova et Round | 0 | 0 | 0 | 0 |
| PKUE | *Psammothidium kuelbsii* | (Lange-Bertalo t in L.-B. & K.) Bukhtiyarova et Round | 0 | 0 | 0 | 0 |
| PLFR | *Planothidium frequentissim um* | (Lange-Bertalo t) Lange-Bertalot | 0 | 0 | 0 | 0 |
| PLFR | *Planothidium frequentissim um* | (Lange-Bertalo t) Lange-Bertalot | 0 | 0 | 0 | 0 |
| PLFR | *Planothidium frequentissim um* | (Lange-Bertalo t) Lange-Bertalot | 0 | 0 | 0 | 0 |
| PLHO | *Platessa holsatica* | (Hustedt) Lange-Bertalot | 0 | 0 | 0 | 0 |
| PLHU | *Platessa hustedtii* | (Krasske) Lange-Bertalot | 0 | 0 | 0 | 0 |
| PLJO | *Platessa joursacense* | (Héribaud) Chudaev in Chudaev. Gololobova et Kulikovskiy | 0 | 0 | 0 | 0 |
| PLPM | *Planothidium pumilum* | Bąk & Lange-Bertalot | 0 | 0 | 0 | 0 |
| PLVA | *Psammothidium levanderi* | (Hustedt) Bukhtiyarova | 0 | 0 | 0 | 0 |

| | | | Alert taxa (1: yes ; 0: no) | | | |
|---|---|---|---|---|---|---|
| code | name | author | BOD5 | SP | NKJ | Pt |
| PLVA | *Psammothidium levanderi* | (Hustedt) Czarnecki in Czarnecki et Edlund | 0 | 0 | 0 | 0 |
| PLVU | *Psammothidium lacus-vulcani* | (Lange-Bert. et Kram.) Bukht. et Round | 0 | 0 | 0 | 0 |
| PMNT | *Planothidium minutissimum* | (Krasske) Lange-Bertalot | 0 | 0 | 0 | 0 |
| PMNT | *Planothidium minutissimum* | (Krasske) Morales | 0 | 0 | 0 | 0 |
| PMOC | *Pseudofallacia monoculata* | (Hustedt) Liu Kociolek & Wang | 0 | 0 | 0 | 0 |
| PMSC* | *Pseudostaurosira microstriata var. microstriata* | (Marciniak) Flower | 0 | 0 | 0 | 0 |
| PMUL | *Planothidium minusculum* | (Hustedt) Witkowski, Kulikovskiy et Pliński | 0 | 0 | 0 | 0 |
| POBL | *Platessa oblongella* | (Østrup) C.E. Wetzel, Lange-Bertalot & Ector | 0 | 0 | 0 | 0 |
| POCL | *Pantocsekiella ocellata* | (Pantocsek) K.T. Kiss et Ács | 0 | 0 | 0 | 0 |
| POVA | *Punctastriata ovalis* | Williams & Round | 0 | 0 | 0 | 0 |
| PPRS | *Pseudostaurosira parasitica* | (W.Smith) Morales | 0 | 0 | 0 | 0 |
| PPSA | *Placoneis pseudanglica* | (Lange-Bertalot) Cox | 0 | 1 | 0 | 0 |
| PRBU | *Planothidium robustius* | (Hustedt) Lange-Bertalot | 0 | 0 | 0 | 0 |
| PROH | *Planothidium rostratoholarcticum* | Lange-Bertalot & Bąk | 1 | 1 | 1 | 0 |
| PROS | *Psammothidium rossii* | (Hustedt) Bukhtiyarova et Round | 0 | 0 | 0 | 0 |
| PRST | *Planothidium rostratum* | (Østrup) Lange-Bertalot | 0 | 1 | 0 | 0 |
| PRST | *Planothidium rostratum* | (Østrup) Round & Bukhtiyarova | 0 | 1 | 0 | 0 |
| PSBR | *Pseudostaurosira brevistriata* | (Grun.in Van Heurck) Williams & | 0 | 0 | 0 | 0 |

| code | name | author | Alert taxa (1: yes ; O: no) BOD5 | SP | NKJ | Pt |
|---|---|---|---|---|---|---|
| | | Round | | | | |
| PSCA* | *Pinnularia subcapitata var. subcapitata* | Gregory | 0 | 0 | 0 | 0 |
| PSCT | *Psammothidium scoticum* | (Flower & Jones) Bukhtiyarova et Round | 0 | 0 | 0 | 0 |
| PSME | *Pseudostauros ira medliniae* | D.M.Williams & Morales | 0 | 0 | 0 | 0 |
| PSPO | *Pseudostauros ira polonica* | (Witak & Lange-Bertalot ) Morales et M.B. Edlund | 0 | 0 | 0 | 0 |
| PSRE | *Psammothidium rechtensis* | (Leclercq) Lange-Bertalot | 0 | 0 | 0 | 0 |
| PSSE | *Pseudostauros ira elliptica* | (Gasse) Jung & Medlin | 0 | 0 | 0 | 0 |
| PSSE | *Pseudostauros ira elliptica* | (Schumann) Edlund, Morales & Spaulding | 0 | 0 | 0 | 0 |
| PSYM | *Placoneis symmetrica* | (Hustedt) Lange-Bertalot | 0 | 0 | 0 | 0 |
| PTCO | *Platessa conspicua* | (A.Mayer) Lange-Bertalot | 0 | 0 | 0 | 0 |
| PTDE | *Planothidium delicatulum* | (Kütz.) Round & Bukhtiyarova | 0 | 1 | 0 | 1 |
| PTDU | *Planothidium dubium* | (Grunow) Round & Bukhtiyarova | 0 | 0 | 0 | 0 |
| PTLA | *Planothidium lanceolatum* | (Brébisson) Round et Bukhtiyarova | 0 | 0 | 0 | 0 |
| PTLA | *Planothidium lanceolatum* | (Brébisson ex Kützing) Lange-Bertalot | 0 | 0 | 0 | 0 |
| PTLA | *Planothidium lanceolatum* | (Brébisson in Kützing) Bukhtiyarova | 0 | 0 | 0 | 0 |
| PTPU | *Praestephanos triporus* | (Genkal & G.V. Kuzmin)Tuji & J.-S. Ki | 1 | 1 | 1 | 1 |
| PUDI | *Punctastriata discoidea* | Flower | 0 | 0 | 0 | 0 |
| PULA | *Punctastriata lancettula* | (Schumann) Hamilton & Siver | 0 | 0 | 0 | 0 |
| PUSB | *Pseudostauros ira* | (Grunow) Kulikovskiy & | 0 | 0 | 0 | 0 |

| code | name | author | Alert taxa (1: yes ; O: no) | | | |
|---|---|---|---|---|---|---|
| | | | BOD5 | SP | NKJ | Pt |
| | *subconstricta* | Genkal ., stat. nov. | | | | |
| PVEN | *Psammothidium ventrale* | (Krasske) Bukhtiyarova et Round | 0 | 0 | 0 | 0 |
| PWUE | *Pantocsekiella wuethrichiana* | (Druart et F. Straub) K.T. Kiss et Ács | 0 | 0 | 0 | 0 |
| PZIE | *Platessa ziegleri* | (Lange-Bertalot) Lange-Bertalot | 0 | 0 | 0 | 0 |
| RABB | *Rhoicosphenia abbreviata* | (C.Agardh) Lange-Bertalot | 1 | 0 | 1 | 1 |
| RFON | *Reimeria fontinalis* | Levkov, & Ector | 0 | 0 | 0 | 0 |
| ROVA | *Reimeria ovata* | (Hustedt) Levkov, & Ector | 0 | 0 | 0 | 0 |
| RPUS | *Rossithidium pusillum* | (Grunow) F.E.Round & Bukhtiyarova | 0 | 0 | 0 | 0 |
| RSIN | *Reimeria sinuata* | (Gregory) Kociolek & Stoermer | 0 | 0 | 0 | 0 |
| RUNI | *Reimeria uniseriata* | Sala Guerrero & Ferrario | 0 | 0 | 0 | 0 |
| SACB | *Sellaphora archibaldii* | (J.C. Taylor et Lange-Bertalot) Ács, C.E. Wetzel et Ector . | 0 | 0 | 0 | 0 |
| SANG | *Surirella angusta* | Kützing | 0 | 0 | 0 | 0 |
| SARV | *Sellaphora arvensis* | (Hustedt) C.E. Wetzel et Ector | 0 | 0 | 0 | 0 |
| SBND | *Staurosira binodis* | (Ehrenberg) Lange-Bertalot in Hofmann Werum et Lange-Bertalot | 0 | 0 | 0 | 0 |
| SBND | *Staurosira binodis* | (Ehrenb.) Kulikovskiy & Genkal | 0 | 0 | 0 | 0 |
| SBOH | *Surirella bohemica* | Maly | 0 | 0 | 0 | 0 |
| SBRE* | *Surirella brebissonii var. brebissonii* | Krammer & Lange-Bertalot | 0 | 0 | 0 | 0 |
| SCAN | *Staurosirella canariensis* | (Lange-Bertalot) E. Morales, | 0 | 0 | 0 | 1 |

| code | name | author | Alert taxa (1: yes ; 0: no) BOD5 | SP | NKJ | Pt |
|---|---|---|---|---|---|---|
| | | Ector, Maidana & Grana . | | | | |
| SCHK | *Sellaphora chistiakovae* | (Kulikovskiy et Lange-Bertalot ) Wetzel, Ector Van De Vijver,Compère & D.G.Mann | 0 | 0 | 0 | 0 |
| SCON | *Staurosira construens* | Ehrenberg | 0 | 0 | 0 | 0 |
| SCPO | *Sellaphora cosmopolitana* | (Lange-Bertalot) C.E. Wetzel et Ector | 0 | 0 | 0 | 0 |
| SCRA | *Sellaphora crassulexigua* | (Reichardt) Wetzel, Ector, Van De Vijver, Compère & D.G.Mann | 1 | 1 | 1 | 1 |
| SCRM | *Stauroneis charlesreimeri* | Lange-Bertalot & Metzeltin | 0 | 0 | 0 | 0 |
| SEAT | *Sellaphora atomoides* | Wetzel & Ector | 0 | 0 | 0 | 0 |
| SEBA | *Sellaphora bacillum* | (Ehrenberg) D.G.Mann | 0 | 0 | 0 | 0 |
| SECA | *Sellaphora capitata* | D.G. Mann & S.M. Mc Donald | 0 | 0 | 0 | 0 |
| SELA | *Sellaphora laevissima* | (Kützing) D.G. Mann | 0 | 0 | 0 | 0 |
| SELO | *Sellaphora elorantana* | (Lange-Bertalot) C.E. Wetzel | 0 | 0 | 0 | 0 |
| SEUT | *Sellaphora utermoehlii* | (Hustedt) C.E. Wetzel et D.G. Mann | 0 | 0 | 0 | 0 |
| SEXG | *Stauroforma exiguiformis* | (Lange-Bertalot) Flower Jones et Round | 0 | 0 | 0 | 0 |
| SGRL | *Stauroneis gracilior* | Reichardt | 0 | 0 | 0 | 0 |
| SGRL | *Stauroneis gracilior* | Reichardt in Van de Vijver & al. | 0 | 0 | 0 | 0 |
| SHAN | *Stephanodiscus hantzschii* | Grunow in Cleve & Grunow | 0 | 0 | 0 | 0 |
| SIDE | *Simonsenia delognei* | Lange-Bertalot | 0 | 1 | 1 | 1 |
| SINM | *Stauroforma inermis* | Flower Jones et Round | 1 | 1 | 0 | 1 |
| SKOS | *Skabitschewsk* | (Cleve-Euler) | 0 | 0 | 0 | 0 |

| code | name | author | Alert taxa (1: yes ; O: no) | | | |
|---|---|---|---|---|---|---|
| | | | BOD5 | SP | NKJ | Pt |
| | *ia œstrupii* | Kulikovskiy & Lange-Bertalot | | | | |
| SKPO | *Skeletonema potamos* | (Weber) Hasle | 0 | 0 | 0 | 0 |
| SKRM | *Staurosirella krammeri* | E.A.Morales, C.Wetzel & Ector | 0 | 0 | 0 | 0 |
| SLAC | *Surirella lacrimula* | English | 0 | 0 | 0 | 0 |
| SLEP | *Staurosirella leptostauron* | (Ehr.) Williams & Round | 0 | 0 | 0 | 0 |
| SLMU | *Staurosirella mutabilis* | (W. Smith) E. Morales & Van de Vijver | 0 | 0 | 0 | 0 |
| SLPP | *Staurosira lapponica* | (Grunow) Lange-Bertalot | 0 | 0 | 0 | 0 |
| SMTO | *Sellaphora mutatoides* | Lange-Bertalot & Metzeltin | 0 | 0 | 0 | 0 |
| SNEO | *Stephanodiscus neoastraea* | Håkansson et Hickel | 0 | 0 | 0 | 0 |
| SNIG | *Sellaphora nigri* | C.E. Wetzel et Ector . emend | 1 | 1 | 1 | 1 |
| SNPI | *Staurosirella neopinnata* | E.A. Morales, C.E. Wetzel, E.Y. Haworth & L. Ector | 0 | 0 | 0 | 0 |
| SODB | *Staurosira oldenburgiana* | (Hustedt)Lange-Bertalot | 0 | 1 | 0 | 1 |
| SPCO | *Staurosira pseudoconstruens* | (Marciniak) Lange-Bertalot | 0 | 0 | 0 | 0 |
| SPDV | *Sellaphora pseudoarvensis* | (Hustedt) C.E. Wetzel et Ector | 0 | 0 | 0 | 1 |
| SPHO | *Stauroneis phoenicenteron* | (Nitzsch.) Ehrenberg | 0 | 0 | 0 | 0 |
| SPIN | *Staurosirella pinnata* | (Ehrenberg) Williams&Round | 0 | 0 | 0 | 0 |
| SPRG | *Skabitschewskia peragalli* | (Brun & Héribaud) Kulikovskiy & Lange-Bertalot | 0 | 0 | 0 | 0 |
| SPSV | *Sellaphora pseudoventralis* | (Hustedt) Chudaev et Gololobova | 0 | 0 | 0 | 0 |
| SPSV | *Sellaphora pseudoventralis* | (Hustedt) Wetzel, Ector Van De Vijver, | 0 | 0 | 0 | 0 |

| code | name | author | Alert taxa (1: yes ; O: no) | | | |
|---|---|---|---|---|---|---|
| | | | BOD5 | SP | NKJ | Pt |
| | | Compère & D.G.Mann. Mann | | | | |
| SPUP | *Sellaphora pupula* | (Kützing) Mereschkowksy | 1 | 1 | 1 | 0 |
| SRAE | *Sellaphora raederae* | (Lange-Bertalot) C.E. Wetzel | 0 | 0 | 0 | 0 |
| SRBU | *Staurosira robusta* | (Fusey) Lange-Bertalot | 0 | 0 | 0 | 0 |
| SRHE | *Sellaphora rhombelliptica* | (Gerd Moser, Lange-Bertalot et Metzeltin) C.E. Wetzel et Ector | 0 | 0 | 0 | 0 |
| SRMA | *Staurosira martyi* | (Héribaud) Lange-Bertalot | 0 | 0 | 0 | 0 |
| SRTU | *Sellaphora rotunda* | (Hustedt) Wetzel, Ector Van De Vijver, Compère & D.G.Mann. Mann | 0 | 0 | 0 | 0 |
| SSBG | *Sellaphora schaumburgii* | (Lange-Bertalot et G. Hofmann) C.E. Wetzel & Ector | 0 | 0 | 0 | 0 |
| SSGE | *Sellaphora saugerresii* | (Desm.) C.E. Wetzel & D.G. Mann in Wetzel et al. | 1 | 1 | 1 | 1 |
| SSMI | *Stauroneis smithii* | Grunow | 0 | 0 | 0 | 0 |
| SSRT | *Sellaphora subrotundata* | (Hustedt) Wetzel, Ector Van De Vijver, Compère & D.G.Mann. Mann | 0 | 0 | 0 | 0 |
| SSTM | *Sellaphora stroemii* | (Hustedt) Kobayasi in Mayama Idei Osada & Nagumo | 0 | 0 | 0 | 0 |
| SSVE | *Staurosira venter* | (Ehrenberg) Cleve & Moeller | 0 | 0 | 0 | 0 |
| SSVE | *Staurosira venter* | (Ehr.) H.Kobayasi | 0 | 0 | 0 | 0 |
| SSVE | *Staurosira venter* | (Ehrenberg) Grunow in Pantocsek | 0 | 0 | 0 | 0 |
| STLG | *Staurosirella grunowii* | (Pantocsek) E. Morales, Buczkó & Ector | 0 | 0 | 0 | 0 |
| STMI | *Stephanodiscu* | (Kützing) | 0 | 0 | 0 | 0 |

| code | name | author | Alert taxa (1: yes ; O: no) BOD5 | SP | NKJ | Pt |
|---|---|---|---|---|---|---|
| | *s minutulus* | Cleve & Moller | | | | |
| STOV | *Staurosirella ovata* | Morales | 0 | 0 | 0 | 0 |
| STSB | *Staurosira berolinensis* | (Lemm.)Kulikovskiy & Genkal | 0 | 0 | 0 | 0 |
| STSB | *Staurosira berolinensis* | (Lemm.) Lange-Bertalot | 0 | 0 | 0 | 0 |
| STSE | *Stauroneis separanda* | Lange-Bertalot & Werum | 0 | 0 | 0 | 0 |
| STTU | *Stephanodiscus tenuis* | Hustedt | 0 | 0 | 0 | 0 |
| SULI | *Surirella librile* | (Ehrenberg Ehrenberg | 0 | 0 | 0 | 0 |
| SUUN | *Surirella undulata* | (Ehrenberg) Ehrenberg | 0 | 0 | 0 | 0 |
| SVTB | *Sellaphora vitabundicta* | E. Reichardt nov. spec. | 0 | 0 | 0 | 0 |
| SVTL | *Sellaphora ventraloides* | (Hustedt) Falasco & Ector | 0 | 0 | 0 | 0 |
| TANG | *Tryblionella angustata* | W.M. Smith | 0 | 0 | 0 | 0 |
| TATU | *Tryblionella angustatula* | (Lange-Bertalot) Cantonati & Lange-Bertalot in Kusber et al. . | 0 | 1 | 0 | 1 |
| TBNO | *Tryblionella brunoi* | (Lange-Bertalot) Cantonati et Lange-Bertalot in Kusber et al. | 0 | 0 | 0 | 0 |
| TCAL | *Tryblionella calida* | (Grunow in Cl. & Grun.) D.G. Mann in Round Crawford & Mann | 0 | 0 | 0 | 0 |
| TFAS | *Tabularia fasciculata* | (Agardh)Williams et Round | 0 | 0 | 0 | 0 |
| TFEN | *Tabellaria fenestrata* | (Lyngbye) Kützing | 0 | 0 | 0 | 0 |
| TFLO | *Tabellaria flocculosa* | (Roth) Kützing | 0 | 0 | 0 | 0 |
| TKUE | *Tryblionella kuetzingii* | Alvarez-Blanco & S.Blanco | 0 | 0 | 0 | 0 |
| TLEV | *Tryblionella levidensis* | Wm. Smith | 0 | 0 | 0 | 0 |
| TVEN | *Tabellaria ventricosa* | Kützing | 0 | 0 | 0 | 0 |
| UACU | *Ulnaria acus* | (Kützing) Aboal | 1 | 1 | 1 | 1 |

| | | | Alert taxa (1: yes ; O: no) | | | |
|---|---|---|---|---|---|---|
| code | name | author | BOD5 | SP | NKJ | Pt |
| UBIC | *Ulnaria biceps* | (Kützing) Compère | 0 | 0 | 0 | 0 |
| UDEL | *Ulnaria delicatissima* | (W.Smith) Aboal & Silva | 0 | 0 | 0 | 0 |
| UDEL | *Ulnaria delicatissima* | (W.Smith) Aboal | 0 | 0 | 0 | 0 |
| UGRU | *Ulnaria grunowii* | (Lange-Bertalot et Ulrich) Cantonati et Lange-Bertalot in Kusber & al. | 0 | 0 | 0 | 0 |
| UULN | *Ulnaria ulna* | (Nitzsch) Compère | 1 | 0 | 1 | 1 |
| VUCO | *Diatomées non identifiées vc* | non identifiées vue connectives | 0 | 0 | 0 | 0 |
| ZZZZ | *Genre non identifie* | NA | 0 | 0 | 0 | 0 |
