## Supplementary material 1 Table 2 for "A new diatom-based multimetric index to assess lake ecological status"

| variable | type | substrate | Mgroup | SDgroup |
|---|---|---|---|---|
| Cond | LA | mineral | 0.98 | 0.02 |
| Cond | MA | mineral | 0.87 | 0.16 |
| Cond | HA | mineral | 0.55 | 0.22 |
| Cond | LA | macrophyte | 0.99 | 0.01 |
| Cond | MA | macrophyte | 0.95 | 0.06 |
| Cond | HA | macrophyte | 0.62 | 0.21 |
| BOD5 | LA | mineral | 0.98 | 0.03 |
| BOD5 | MA | mineral | 0.82 | 0.20 |
| BOD5 | HA | mineral | 0.88 | 0.18 |
| BOD5 | LA | macrophyte | 0.99 | 0.02 |
| BOD5 | MA | macrophyte | 0.87 | 0.19 |
| BOD5 | HA | macrophyte | 0.85 | 0.22 |
| MES | LA | mineral | 0.96 | 0.04 |
| MES | MA | mineral | 0.79 | 0.21 |
| MES | HA | mineral | 0.74 | 0.27 |
| MES | LA | macrophyte | 0.98 | 0.02 |
| MES | MA | macrophyte | 0.90 | 0.19 |
| MES | HA | macrophyte | 0.82 | 0.24 |
| NKJ | LA | mineral | 0.97 | 0.03 |
| NKJ | MA | mineral | 0.87 | 0.20 |
| NKJ | HA | mineral | 0.90 | 0.16 |
| NKJ | LA | macrophyte | 0.98 | 0.02 |
| NKJ | MA | macrophyte | 0.91 | 0.18 |
| NKJ | HA | macrophyte | 0.87 | 0.21 |
| NO2 | LA | mineral | 0.96 | 0.06 |
| NO2 | MA | mineral | 0.83 | 0.15 |
| NO2 | HA | mineral | 0.75 | 0.17 |
| NO2 | LA | macrophyte | 0.98 | 0.02 |
| NO2 | MA | macrophyte | 0.84 | 0.20 |
| NO2 | HA | macrophyte | 0.77 | 0.19 |
| NO3 | LA | mineral | 0.89 | 0.11 |
| NO3 | MA | mineral | 0.85 | 0.16 |
| NO3 | HA | mineral | 0.58 | 0.18 |
| NO3 | LA | macrophyte | 0.92 | 0.08 |
| NO3 | MA | macrophyte | 0.90 | 0.13 |
| NO3 | HA | macrophyte | 0.68 | 0.16 |
| O2 | LA | mineral | 0.94 | 0.05 |
| O2 | MA | mineral | 0.94 | 0.08 |
| O2 | HA | mineral | 0.97 | 0.06 |
| O2 | LA | macrophyte | 0.96 | 0.04 |
| O2 | MA | macrophyte | 0.94 | 0.12 |
| O2 | HA | macrophyte | 0.97 | 0.08 |
| PO4 | LA | mineral | 0.97 | 0.03 |
| PO4 | MA | mineral | 0.84 | 0.21 |
| PO4 | HA | mineral | 0.78 | 0.25 |

| variable | type | substrate | Mgroup | SDgroup |
|---|---|---|---|---|
| PO4 | LA | macrophyte | 0.98 | 0.02 |
| PO4 | MA | macrophyte | 0.94 | 0.14 |
| PO4 | HA | macrophyte | 0.84 | 0.23 |
| Pt | LA | mineral | 0.93 | 0.06 |
| Pt | MA | mineral | 0.73 | 0.23 |
| Pt | HA | mineral | 0.79 | 0.25 |
| Pt | LA | macrophyte | 0.98 | 0.02 |
| Pt | MA | macrophyte | 0.83 | 0.24 |
| Pt | HA | macrophyte | 0.85 | 0.21 |
| %O2 | LA | mineral | 0.99 | 0.01 |
| %O2 | MA | mineral | 0.99 | 0.03 |
| %O2 | HA | mineral | 1.00 | 0.01 |
| %O2 | LA | macrophyte | 1.00 | 0.01 |
| %O2 | MA | macrophyte | 0.97 | 0.07 |
| %O2 | HA | macrophyte | 0.99 | 0.03 |
