## Supplementary material 1 Table 1 for "A new diatom-based multimetric index to assess lake ecological status"

| variable | Min | Max |
|---|---|---|
| BOD5 | -4.248347 | 0.900000 |
| MES | -4.158029 | 1.000000 |
| NKJ | -4.804299 | 1.000000 |
| NO2 | -5.388889 | 1.441964 |
| NO3 | -4.191932 | 2.234127 |
| PO4 | -5.271100 | 1.000000 |
| Pt | -3.559219 | 1.173913 |
| cond__invivo | -5.280570 | 1.964286 |
| o2_dissous__invivo | -7.668478 | 1.200000 |
| sat_o2__invivo | -12.871176 | 1.000000 |
